# Interactions between auditory statistics processing and visual experience emerge only in late development

**DOI:** 10.1101/2020.12.16.423049

**Authors:** Martina Berto, Pietro Pietrini, Emiliano Ricciardi, Davide Bottari

## Abstract

The human auditory system relies on both detailed and summarized representations to recognize different sounds. As local features can exceed the storage capacity, average statistics are computed over time to generate more compact representations at the expense of temporal details availability. This study aimed to identify whether these fundamental sound analyses develop and function exclusively under the influence of the auditory system or interact with other modalities, such as vision. We employed a validated computational synthesis approach allowing to control directly statistical properties embedded in sounds. To address whether the two modes of auditory representation (local features processing and statistical averaging) are influenced by the availability of visual input in different phases of development, we tested samples of sighted controls (SC), congenitally blind (CB), and late-onset (> 10 years of age) blind (LB) individuals in two separate experiments which uncovered auditory statistics computations from behavioral performances. In experiment 1, performance relied on the availability of local features at specific time points; in experiment 2, performance benefited from computing average statistics over longer durations. As expected, when sound duration increased, detailed representation gave way to summary statistics in SC. In both experiments, the sample of CB individuals displayed a remarkably similar performance revealing that both local and global auditory processes are not altered by blindness since birth. Conversely, LB individuals performed poorly compared to the other groups when relying on local features, with no impact on statistical averaging. The dampening in the performance was not associated with the onset and duration of visual deprivation. Results provide clear evidence that vision is not necessary for the development of the auditory computations tested here. Remarkably, a functional interplay between acoustic details processing and vision emerges at later developmental phases. Findings are consistent with a model in which the efficiency of local auditory processing is vulnerable in case sight becomes unavailable. Ultimately results are in favor of a shared computational framework for auditory and visual processing of local features, which emerges in late development.

## INTRODUCTION

The auditory system is specialized in capturing fine-grained details from sound waves (Plomp, 1964). These local temporal features are then integrated over time by mechanisms that retain, summarize, and abstract them into more compact acoustic percepts (Yabe et al., 1998; McDermott, Schemitsch, and Simoncelli, 2013). Computational synthesis approaches, derived from information theories (Barlow, 1961), allowed to investigate these processes by implementing mathematical models to describe the set of measurements that the auditory system operates (McDermott and Simoncelli, 2011; McDermott, Schemitsch, and Simoncelli, 2013). Such models have revealed that stationary sounds are analyzed by extracting a set of auditory statistics (McDermott and Simoncelli, 2011) whose processing represents the keystone of acoustic features integration. Auditory statistics result from a set of computations strictly dependent on anatomical and functional properties of the auditory pathway (Ruggero, 1992; Dau, Kollmeier and Kohlrausch, 1997; Gygi, Kidd and Watson, 2004; Joris, Schreiner and Rees, 2004; Baumann et al., 2011), and can be consequently described only through biologically plausible models (McDermott and Simoncelli, 2011). The auditory system processes these statistics by averaging short-term acoustic events (McDermott and Simoncelli, 2011; McDermott, Schemitsch, and Simoncelli, 2013; McWalter and McDermott, 2018), along specific time windows (McWalter and McDermott, 2018) and uses this information to derive compact representations of sound objects. The auditory statistics processing can be broken down into two main modes of representation. (1) Local features processing, by which fine-grained temporal details are extracted from a sound and stored; (2) Statistical averaging, by which local features are averaged within a time-window of integration, resulting in a global representation as local details are no longer retained (McDermott, Schemitsch, and Simoncelli, 2013).

While auditory statistics represent unimodal computations, the auditory system does not develop and function in isolation, and other senses, such as vision, modulate its functional and structural organization. Studies in non-human animal models revealed that, in the early development, the onset of visual input (eyes opening) gates the critical period closure of basic auditory functions. This evidence suggested that visually modulated sensitive periods in the auditory cortex exist (Mowery, Kotak, and Sanes, 2016). In adults, visual events are known to directly modulate auditory neuron responses in both animals (Kayser, Petkov, and Logothetis, 2008) and humans (Thorne et al., 2011). At a functional level, visual systems play an important role in auditory features segregation, helping in disambiguating difficult instances (e.g., Golumbic et al., 2013). A common computational framework for feature extraction might exist between auditory and visual modality (Shamma, 2001). In the same vein, the set of auditory statistics evaluated here were derived by auditory computational models (McDermott and Simoncelli, 2011), which are conceptually very similar to those derived by visual ones (Portilla and Simoncelli, 2000). In the present study, we directly investigated if the auditory statistics development and functioning fall within the exclusive competence of the auditory systems, or are instead influenced by vision.

Visual deprivation models have been systematically employed to indirectly uncover the audio-visual interplay (for review, see Röder, Kekunnaya, and Guerreiro, 2020). The lack of visual inputs has consistently been associated with altered auditory processing across different functions (for review, see Röder and Pavani, 2012), but whether visual input availability alters specific underlying auditory computations is still unknown. In humans, previous evidence has identified visual inputs availability since birth as a prerequisite for the full development of auditory spatial calibration (Gori et al., 2014). On a different note, a large body of evidence suggests that both congenital and late-onset visual deprivation exert a compensating influence over certain higher-order auditory functions (see Röder, Kekunnaya, and Guerreiro, 2020). Compensatory effects have been consistently observed both in the context of spatial processing of auditory stimuli (e.g., Battal et al., 2019) and in tasks requiring spectro-temporal auditory features analyses, such as mnemonic representations of sounds (Röder and Rösler 2003), verbal memory (Amedi et al., 2003), frequency tuning (Huber 2019), speech comprehension (Trouvain, 2007; Dietrich, Hertrich, and Ackermann, 2013), and auditory temporal resolution (Muchnik et al., 1991).

To address whether the two modes of auditory representation, the local features processing and statistical averaging, are differently influenced by visual input availability, we tested blind populations against sighted individuals in two experiments designed to tap into these two computational modes. By combining a sound synthesis algorithm (McDermott and Simoncelli, 2011; McDermott, Schemitsch, and Simoncelli, 2013) with psychophysics, the methodology adopted here provided the opportunity to uncover local and global auditory statistics computations from the discriminative responses elicited by different sound properties. Importantly, comparing samples of individuals who were visually deprived since birth or in late phases of development allowed to assess whether the selected aspect of sound representation interacts with vision at specific time points along the life span.

We could expect detrimental behavioral effects associated with the lack of vision (e.g., Gori et al., 2014) in one of the samples of blind individuals or both. This outcome would syndicate for a modality interplay between vision and auditory computations in which vision plays a crucial role in supporting specific aspects of basic auditory processing (i.e., acoustic features segregation; Park et al., 2016). Noticeably, some auditory fundamentals, such as periodicity pitch, are innate and not influenced by early experience (Montgomery and Clarkson, 1997). Thus, sensory experience might also not be required for the full development of the functions tested here, but modality interplays could still occur later. However, provided that several auditory processes can benefit from the lack of vision, behavioral compensatory effects in one or both visual deprivation models (congenital and late-onset blindness) could also be expected in our study. Such results would indicate that vision is not necessary for their development or functioning and would provide evidence that a functional adaptation to lack of vision occurs not only for high-order functions but also for selective basic auditory computations. In both scenarios, a difference between CB and LB group would characterize the developmental trajectory of audio-visual interplay in the context of local and global auditory statistics processing.

## RESULTS

We took advantage of an already corroborated methodological pipeline (McDermott, Schemitsch, and Simoncelli, 2013), and produced different synthetic textures built upon original recordings. Thanks to their properties, a specific category of natural sounds, namely Sound Textures, was used (some examples are the rain, fire, bulldozer, typewriting, waterfall sounds; McDermott & Simoncelli, 2011; Saint-Arnaud & Popat, 2006; Schwarz, 2011). These sounds are rich, ubiquitous, and constant over time (McDermott, Schemitsch, and Simoncelli, 2013; McWalter and McDermott, 2018). We selected 54 environmental recordings of Sound Textures (Table S1, Supplementary Information), among the original set implemented by McDermott, Schemitsch, and Simoncelli, 2013. Synthetic stimuli were created using an Auditory Texture Model (McDermott and Simoncelli, 2011), which efficiently simulates computations performed at peripheral stages of the auditory processing. By computing time-averages of non-linear functions, it was possible to measure a set of statistics: envelope marginal moments, reflecting distribution and sparsity of the signal (Lorenzi et al., 1999), envelope cross-correlation between cochlear envelopes, accounting for the presence of broadband events within the signal (McDermott and Simoncelli, 2011), the modulation bands power, providing information about the temporal structure within cochlear channels (Bacon and Wesley Grantham, 1989; Dau, Kollmeier and Kohlrausch, 1997; McDermott and Simoncelli, 2011), and their correlations (McDermott and Simoncelli, 2011). Although the model represents a mathematical approximation, cochlear envelope statistics convey most of the perceptually relevant information about the sound (Smith, Delgutte and Oxenham, 2002; Gygi, Kidd and Watson, 2004; McDermott and Simoncelli, 2011). By using the full set of statistics to generate synthetic sounds, it is possible to obtain compelling exemplars of the same original Sound Texture (McDermott and Simoncelli, 2011). Our experimental stimuli were created accordingly. By imposing the statistics mentioned above on four different white noise samples, we synthesized four different exemplars for each original Sound Texture. Among themselves, synthetic exemplars of the same texture varied only for their local features, while their long-term average statistics matched the sound they were derived from. In other terms, this process resulted in four synthetic sounds whose properties were constrained only by the selected average statistics extracted from the original recording (Figure 1A). This provided us with the unique opportunity to test for specific computations within the auditory statistics processing, thanks to the unparalleled level of control exerted over the statistics present in the synthesized sounds (for details about synthesis procedure, see Materials and Methods).

**Figure 1.**
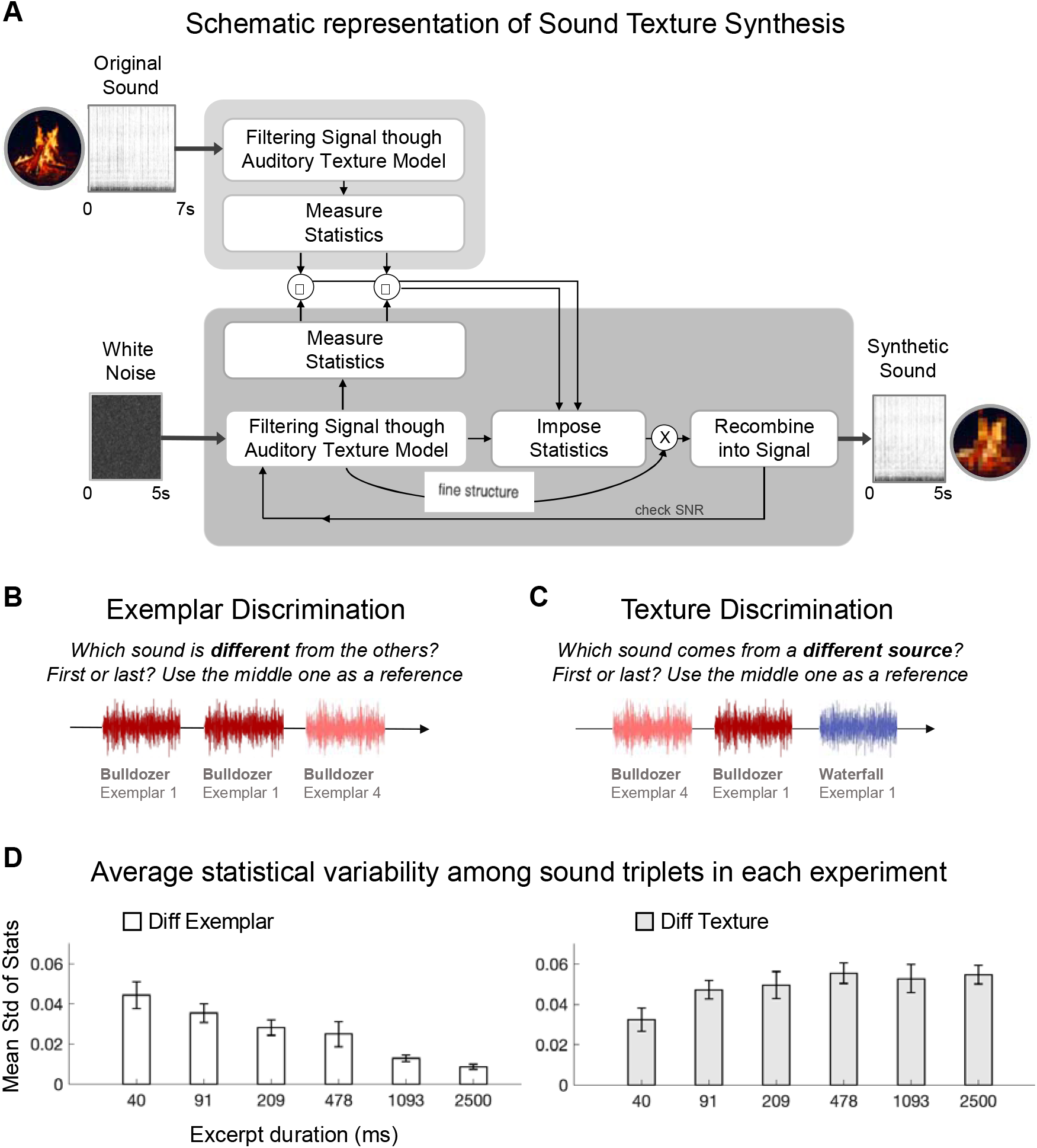
Sound Texture Synthesis, Auditory Statistics Variability, and Experimental Design. (A) Schematic representation of the Sound Texture Synthesis procedure employed to produce synthetic stimuli. A 7-s Original Recording was passed through the Auditory Texture Model (McDermott and Simoncelli, 2011) to extract average statistics of interest. A white noise sample was then passed again through the model while the Original Recording statistics were imposed on its cochlear envelopes. Modified envelopes were multiplied by their associated fine structure and recombined into the signal. This procedure was iterated until a desirable signal-to-noise ratio is reached. The outcome was a Synthetic Sound Texture Exemplar, a 5-s signal containing statistics that matched the original values and was only constrained by them. Adapted from McDermott and Simoncelli, 2011. (B) Exemplar Discrimination. Schematic representation of one trial. Participants had to detect the different sound among the three, being the other two identical. In this case, the correct answer is the last sound since it is another exemplar of the “Bulldozer” Sound Texture. (C) Texture Discrimination. Schematic representation of one trial. Participants had to report which sound was coming from a different source with respect to the other two. In this example, the correct answer is the third sound, being it an excerpt of a different Sound Texture (D) Average variability across a set of statistics (envelope mean, skewness, variance, cross-correlation and modulation power) was computed from couples of excerpts, to measure, in both experiments, the objective statistical difference between reference and deviant sounds. Left: average standard deviation (std) across statistics measured in excerpt pairs originated from Different Exemplars of the same Sound Texture (for all textures in column 1, Table S1, Supplementary Information). When duration increased, average statistics converged to the imposed original values, and variability progressively tended to zero, increasing discrimination difficulty in Exemplar Discrimination. Right: average std across statistics measured in excerpts pairs derived from Different Sound Textures (columns 1 and 2, Table S1, Supplementary Information). In Texture Discrimination, both scenarios (Different Exemplar and Different Texture) were presented and compared against each other. As duration increased, the difference between the two contexts increased, facilitating the recognition of the deviant one. (See also Figure S1 and Table S1)

We tested sighted and blind participants in two selected experiments which included our synthetic sounds and exploited the two modes of auditory statistics representation.

Blind participants were grouped according to blindness onset, whether from birth or developmentally later (>10 years old; see Table 1). Thus, three groups of participants were recruited: congenitally blind (CB), late-onset blind (LB), and sighted control (SC) individuals. All three groups were matched by sample size, age, and gender (see Materials and Methods).

**Table 1.**
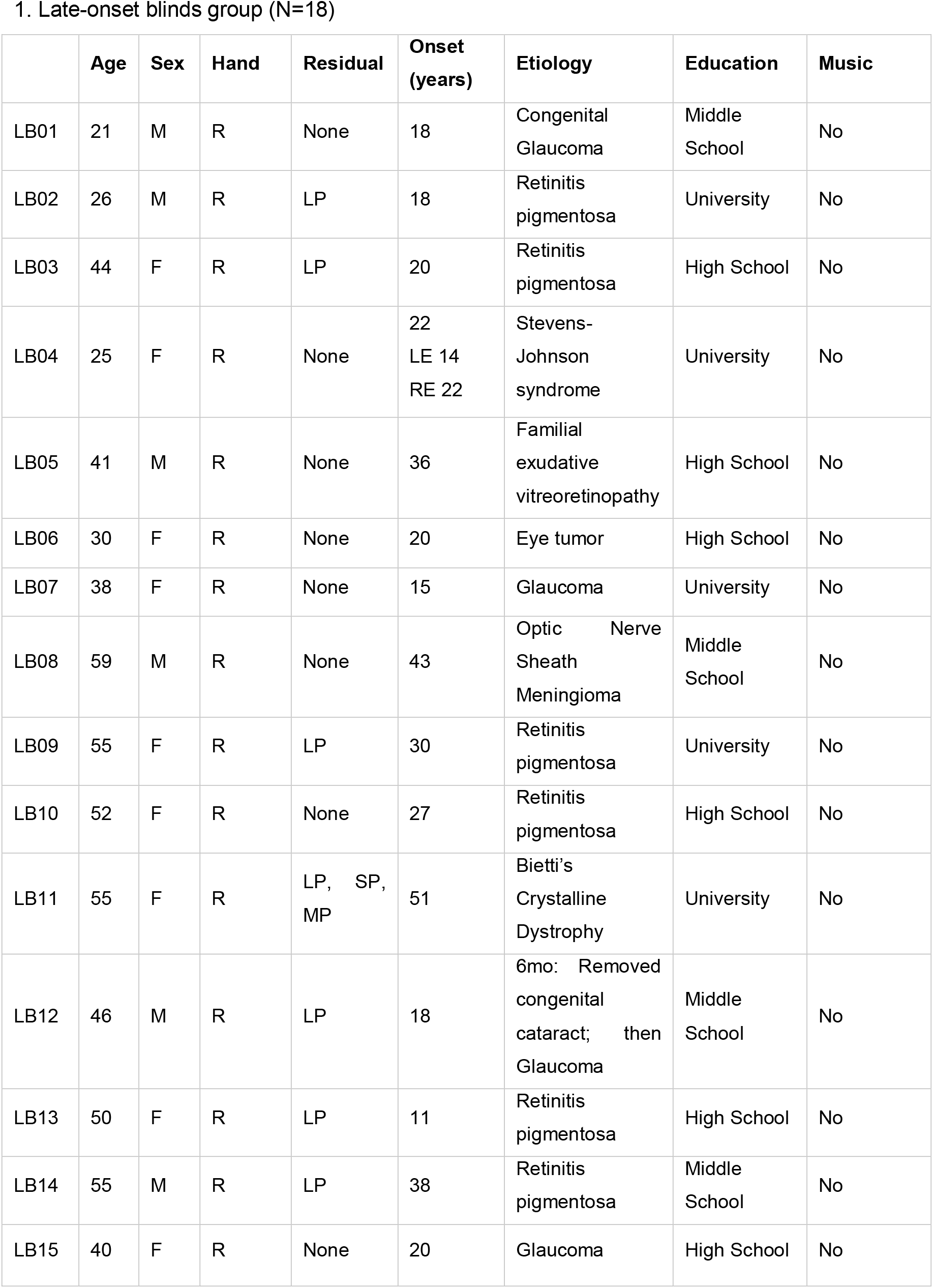
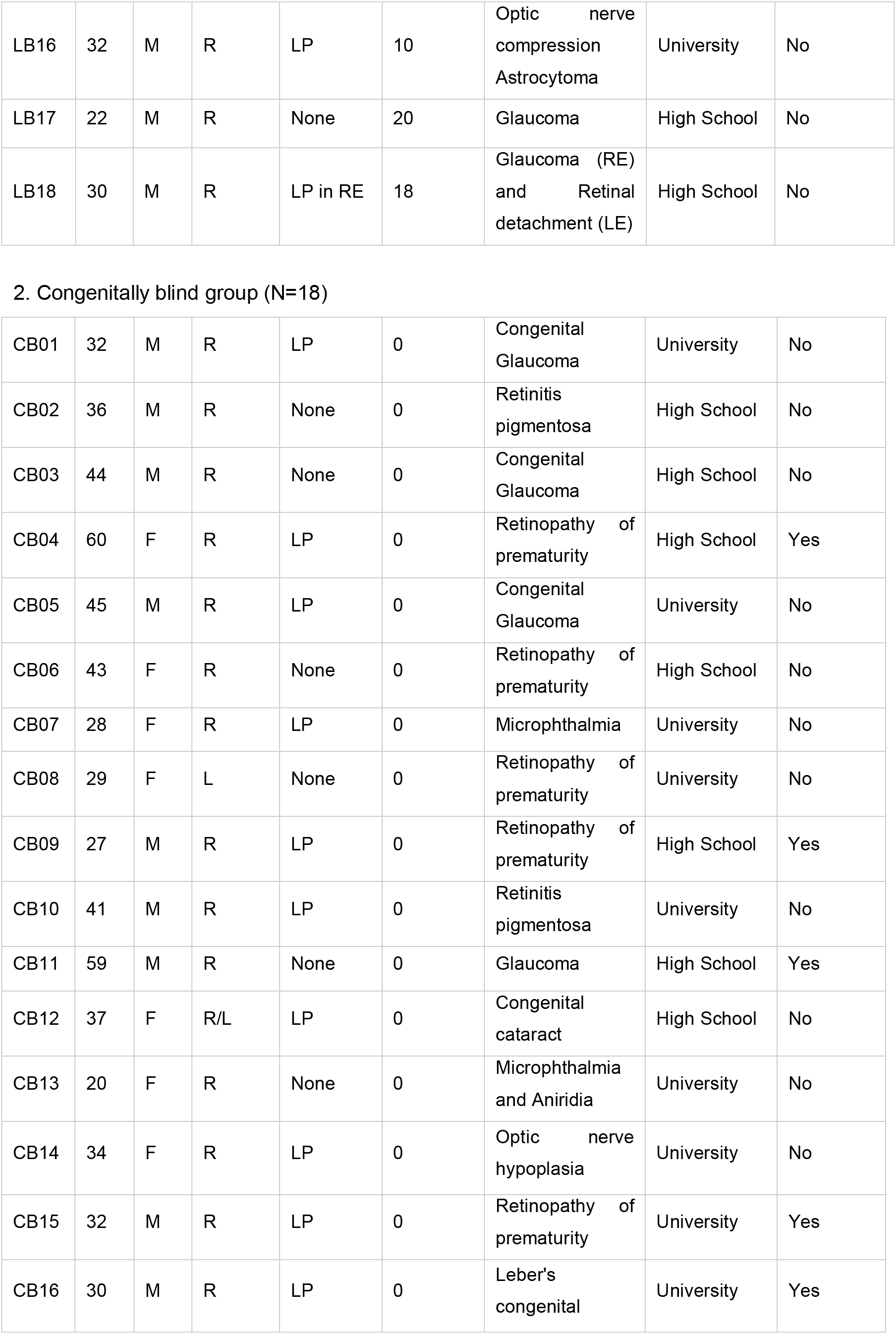
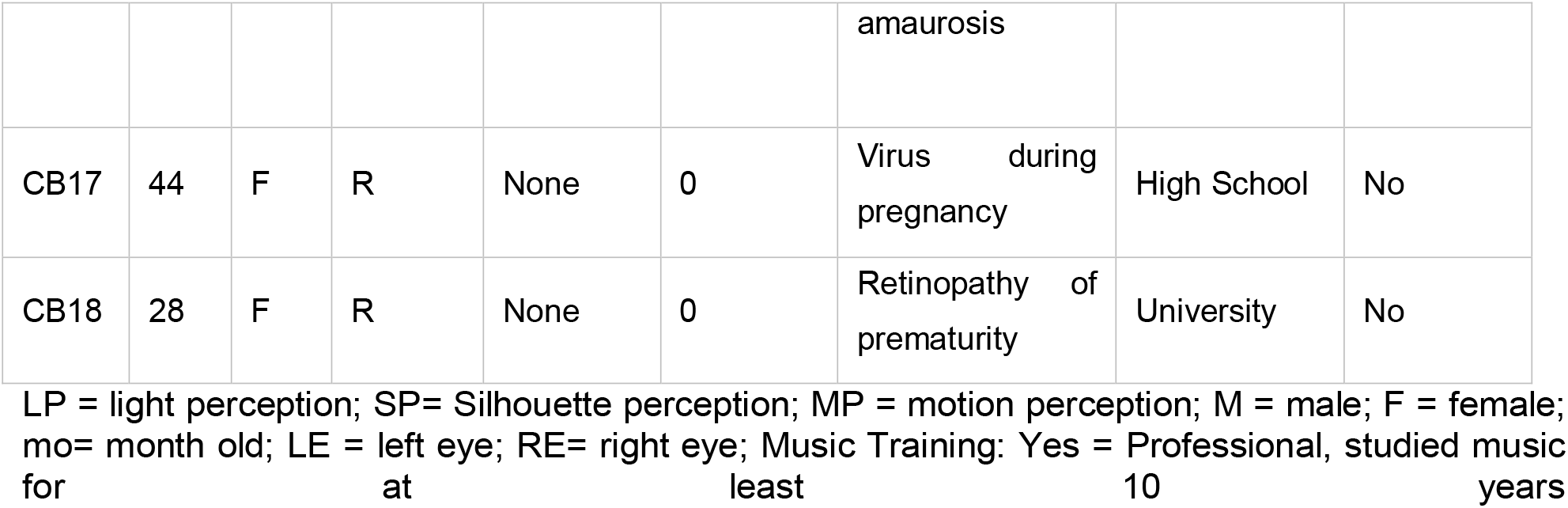
Characteristics of blind participants.

Both experiments consisted of a two-alternative forced-choice oddity (2AFC); participants had to detect the deviant sound among three acoustic samples, selecting between first and third intervals. Stimuli were created by cutting each synthetic exemplar into smaller excerpts of different lengths. Each trial included three excerpts of the same length. The duration of the excerpts varied across trials, for a total of six durations (McDermott, Schemitsch, and Simoncelli, 2013).

In the first experiment, Exemplar Discrimination, participants were asked to report which sound (the first or the third) was different from the other two. Two excerpts were the exact same sound. By contrast, the odd one was an excerpt extracted from a different synthetic exemplar of the same Sound Texture (Figure 1B). Thus, all of them stemmed from the same sound source (e.g., bulldozer), but one was generated starting from a different white noise and consistently diverged for the local features it encompassed as compared to the other two. Nonetheless, as duration increased, average statistics tended to converge towards the original imposed values (Figure 1D, left panel), making all three sounds perceptually very similar and challenging task performance.

In the second experiment, Texture Discrimination, participants were asked to report which sound came from a different acoustic source. Two excerpts were extracted from two distinct synthetic exemplars of the same Sound Texture (e.g., bulldozer), while the deviant was drawn from a synthetic exemplar derived from a different one (e.g., waterfall). Therefore, only the deviant in the triplet comprised both different local features and imposed average statistics (Figure 1C). Since all three sounds represented different excerpts, the local variability between couples was never zero. Still, the average statistics variability between the two sounds originated from the same texture tended to progressively decrease with duration, allowing for the different sound source (the deviant) to emerge perceptually (Figure 1D, right panel).

The employment of these experiments allowed for specific predictions on the outcomes. Consistent with previous findings by McDermott, Schemitsch, and Simoncelli (2013), opposite patterns of results were expected in the two experiments as a function of excerpts duration. For short stimuli, due to differences in features variability among the three sounds, good performance was expected in Exemplar Discrimination compared to Texture Discrimination. By contrast, for long stimuli, statistical averaging was expected to boost Texture Discrimination performance and to hamper Exemplar Discrimination one (McDermott, Schemitsch, and Simoncelli, 2013). Any significant deviation from these expected outcomes indicates changes in the processing of local features or statistical averaging.

The attended pattern of results (McDermott, Schemitsch, and Simoncelli, 2013) was confirmed in our SC group. The performance was better for short durations (40, 91ms) in Exemplar Discrimination as compared to Texture Discrimination (all p < 0.003, corrected). Conversely, for long durations (478, 1093, 2500ms), participants’ accuracy was higher in Texture Discrimination as compared to Exemplar Discrimination (all p < 0.03, corrected). No difference was observed for trials comprising stimuli that were 209ms long (p = 0.31, corrected; Figure 2A). Overall, the data replicated previous findings in sighted individuals, despite participants being blindfolded. Thus, these results represent a validated context to assess whether visual experience impacts auditory statistics processing.

**Figure 2.**
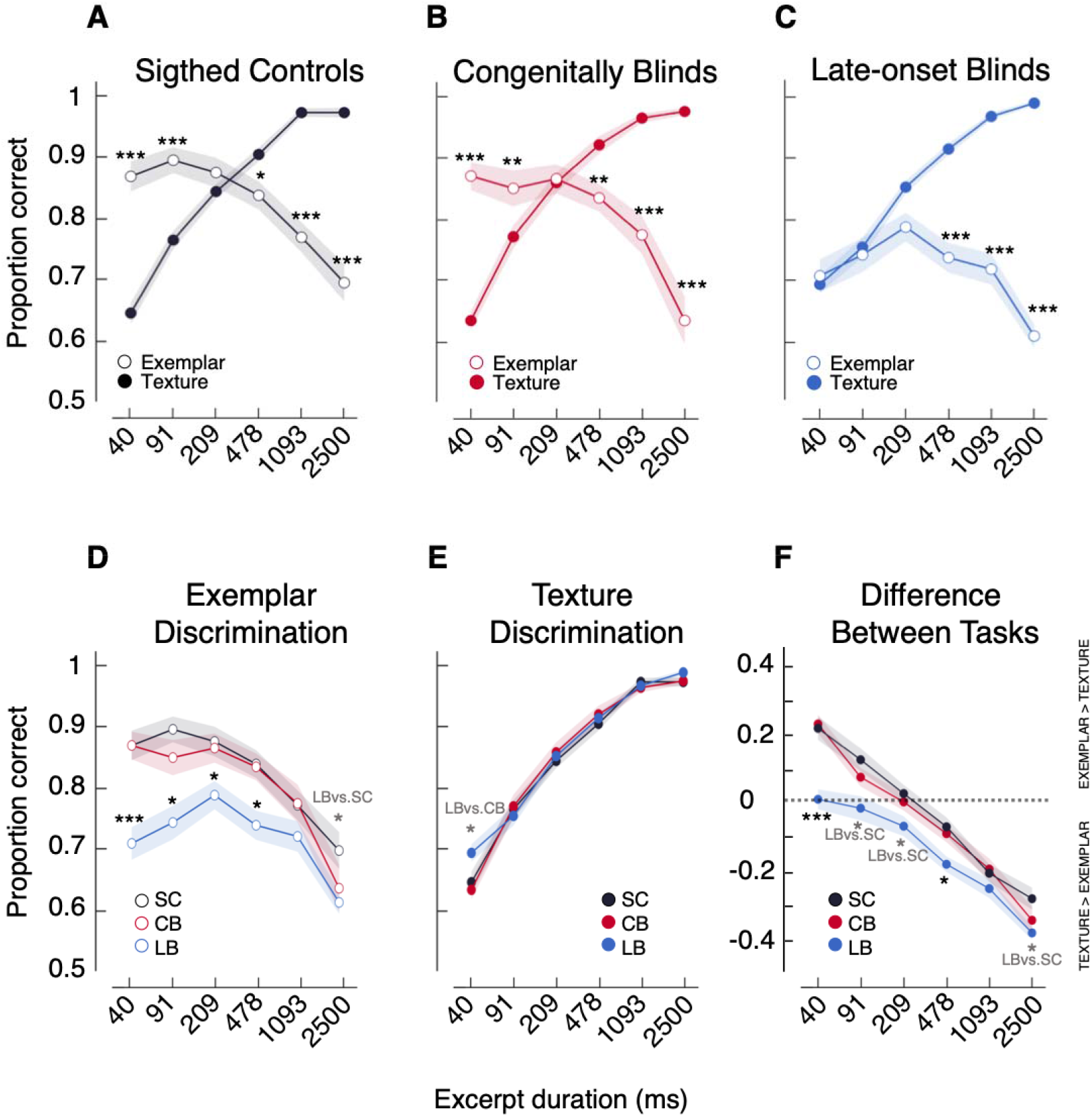
Proportion of correct answers in the two Experiments. (A, B, C) Results for Exemplar Discrimination vs. Texture Discrimination in each group. Proportion of correct answers across individuals at the group level are shown as a function of excerpt duration. (D) Between Group comparisons in Exemplar Discrimination. SC and CB groups’ performance did not differ. The LB group showed an impaired performance compared to the other groups: late sight loss has a detrimental impact when performance could benefit from higher local features variability. (E) Between Group comparisons in Texture Discrimination. No significant differences were observed between SC and CB groups. LB performed better for 40ms stimuli compared to CB. (F) Relative difference between experiments for all three groups. For each group and duration, the averages across participants of the differences between the performance in the two tasks are displayed. Positive values are found when performance in Exemplar is better than in Texture Discrimination, while negative values represent better performance in Texture than Exemplar Discrimination. Unlike the other two groups, LB never showed an advantage in Exemplar Discrimination compared to Texture Discrimination. Shaded regions show interpolated standard error of the mean (SE) at each point. Results were corrected for multiple comparisons using the false discovery rate (FDR). If more than one comparison is significant, stars refer to the lower bound; gray stars indicate that only the labeled comparisons were significant; *** p<.001; ** p<.01; * p<.05. (See also Figure S4).

The CB group displayed a remarkably similar pattern of results. Better performance was found in Exemplar Discrimination for short durations (40, 91ms) compared to Texture Discrimination (all p < 0.02, corrected), and, conversely for long durations (478, 1093, 2500ms) in Texture Discrimination compared to Exemplar (all p < 0.006, corrected). As observed in SC, there was no difference between the two experiments for stimuli that were 209ms long (p = 0.86, corrected; Figure 2B).

By contrast, in the LB group, no significant differences were found between Exemplar Discrimination and Texture Discrimination for all short stimuli (40, 91, 209ms; all p > 0.05, corrected). However, for long durations (478, 1093, 2500ms), LB participants performed better in Texture Discrimination as compared to Exemplar Discrimination (all adjusted p < 0.001), in line with what was observed in the other groups (Figure 2C).

### Late-onset sight loss hampers the processing of local sound features

When contrasting the performance of the SC group with the one of the CB group, no significant differences could be detected across all durations (all p > 0.29, corrected). Results clearly revealed that the processing of fine-grained temporal details is resilient to the absence of visual input since birth.

Conversely, the LB group performance was impaired for almost all of the tested durations as compared to both SC group (40, 91, 209, 478, 2500ms; all p < 0.03, corrected) and CB group (40, 91, 209, 478ms; all p < 0.001, corrected).

Results suggested an altered capability of late blind participants to discriminate sounds when the most efficient strategy is to base performance on local features. These findings reveal that visual input in early phases of development is not a prerequisite to acquiring the ability to discriminate sounds according to their local features. However, this process is encumbered by late-onset blindness (Figure 2D).

### Visual deprivation does not impact statistical averaging

By comparing the performances in Texture Discrimination among the three groups, it was possible to address the impact of visual experience on statistical averaging efficiency.

The performance of the SC group for any of the selected duration (40, 91, 209, 478, 1093, 2500ms) did not differ from both the CB group (all p > 0.47, corrected) and the LB group (all p > 0.06, corrected). When comparing the accuracy of the CB group with the one of the LB group, LB performed better for trials where stimuli were 40ms long (adjusted p = 0.01). No other difference was observed for all the remaining durations (91, 209, 478, 1093, 2500ms; all p > 0.09, corrected; Figure 2E). Altogether, these results reveal that visual deprivation does not significantly influence the process by which auditory statistics are computed over time and used to identify different sound sources.

### Relative difference between experiments reveals no advantages for LB when local features should support the performance

For each participant, the accuracy scores in Texture Discrimination were subtracted from the ones in Exemplar Discrimination, separately for each duration, to compute the relative mean difference between Experiments (Figure 2F). Results showed that compared to both SC and CB groups, this difference was significantly smaller for very short durations in LB group (LC vs. SC: 40, 91; all p < 0.02, corrected; LC vs. CB: 40 p < 0.001, corrected, 91 p = 0.04, uncorrected) and larger for longer ones (LC vs. SC: 209, 478, 2500; all p < 0.03, corrected; LC vs. CB: 478 p < 0.03, corrected; see Figure 2F). Conversely, the relative difference of performance in the two Experiments between CB and SC did not differ for any of the durations (all p > 0.32, corrected; Figure 2F)

These results showed that both CB and SC performed according to predictions showing specifi advantages in one experiment or the other according to stimulus duration. On the contrary, the LB group did not display a boost for those conditions in Exemplar Discrimination in which local feature processing could have helped the performance.

### Ruling out possible confounds

It could be argued that results in the LB group were idiosyncratic of the participants included in the study. However, the sample size was adequate (N = 18 per group) and relatively large compared to previous investigations on blind individuals. All groups were matched adequately by several main variables (size, age, and gender); all participants had no documented auditory deficits, cognitive impairments, or neurological disorders (blindness was associated only to peripheral damage; Table 1). Moreover, LB participants’ performance was hampered only in one of the two experiments. In contrast, the observed results in Texture Discrimination were indistinguishable from the ones of the remaining groups for most of the selected durations. If anything, LB group performance was significantly more efficient at 40ms as compared to CB and partially SC (p < 0.04, uncorrected). In light of these observations, it was possible to exclude that a general cognitive or acoustic impairment could explain the lack of advantage specific for Exemplar Discrimination.

To rule out other possible confounds, we ran specific tests on our sample data. Providing the temporal presentation of triplets in the employed 2AFC protocol, it could have been possible that a particular disposition bias towards the first or last interval was present and diverged among groups, at least partially accounting for different results. This was not the case, as all groups did not significantly differ in terms of predisposition towards stimulus intervals (Figure S2A).

Another possible confounding effect could have been a between-group difference associated with learning or tiredness within an experimental session. It could have been possible that LB group performance had changed as the experiment proceeded (e.g., progressively diminishing across runs, specifically in the Exemplar Discrimination). However, this was not the case. Comparing accuracy across runs, a similar trend in all groups emerged for each duration (Figure 2D).

## DISCUSSION

The present study addressed whether visual experience at different phases of the lifespan exerts a cross-modal influence over specific basic auditory computations. No difference was observed between CB and SC groups. Conversely, we found that LB’s performance was selectively impaired compared to both SC and CB, for those conditions in which optimal performance relied on local features. These results provided evidence that visual experience is not a prerequisite for the full emergence of auditory statistics. Sight loss can negatively impact specific computations only when vision was available in early development and went missing afterward. This outcome is in line with a model in which, in individuals with typical development, visual interactions support auditory segregation in challenging instances accounting for reorganizations and detrimental effects only when visual input goes missing.

### Auditory statistics processing develops regardless of visual input availability

Blindness represents a natural state in which it is possible to address cross-modal interdependencies of sensory functions. By comparing CB with SC groups, our study investigated on one hand whether visual input was necessary for the typical development of the two auditory modes of representation underlying the auditory statistics processing, and on the other hand, whether compensatory mechanisms emerged due to lack of vision since birth. Therefore, two different outcomes could have been expected.

First, if visual experiences were necessary to properly develop the ability to process local features and/or to compute average statistics, we could have observed an impaired performance of CB participants in one of the two experiments, or both, compared to sighted individuals. For example, evidence exists that CB individuals fail at performing auditory space bisection tasks (Gori et al., 2014), and this observation provides evidence that visual experience is necessary to efficiently encode Euclidian coordinates of sounds (Gori et al., 2014; Gori, Amadeo, and Campus, 2020a,b). Our results provide strong evidence for the relative independence of the development of auditory statistic processing from early visual experience.

On the other hand, coherently with previous evidence of enhanced auditory skills in congenitally and early blinds (e.g., Doucet et al., 2005; Muchnik et al., 1991; Röder et al., 1999), lack of vision since birth could have improved the sensitivity to the auditory information included in sound textures, leading to the emergence of specific compensatory mechanisms. For instance, scattered evidence suggests that blind individuals are able to understand syllables at a greater time speed compared to natural speech (Moos and Trouvain, 2007; Trouvain, 2007; Dietrich, Hertrich and Ackermann, 2013), an ability perhaps supported by a higher auditory sampling rate capacity. We could have expected the CB group to be able to retain more local features in time before averaging statistics, showing a better performance for longer durations in Exemplar Discrimination. A compensatory mechanism favoring the statistical averaging mode would have led to better results in Texture Discrimination. Furthermore, a better frequency tuning (Huber et al., 2019) to temporal local features in CB would have enhanced the sensitivity to small acoustic differences, leading to superior-to-sighted performances for short excerpts both in Exemplar and Texture Discrimination. However, none of these hypotheses was confirmed in our data, as the performance was indistinguishable between CB and SC groups. The absence of compensatory mechanisms -arising due to the lack of vision since birth-suggests these basic auditory computations cannot be improved by cross-modal adaptations associated to early blindness.

Previous evidence already suggested the existence of auditory functions which develop independently from visual experience since birth (for instance, auditory gap detection threshold; Weaver and Stevens, 2006; pure tone threshold audiometry and acoustic reflex threshold; Starlinger and Niemeyer, 1981). Similarly, it seems that the development of auditory computations tested in this study is not influenced by the availability of other senses.

We can argue that the extraction of the auditory statistics necessary for textures recognition occurs at a very basic level of processing, while functions that showed visual input dependencies are mostly the result of higher-order cortical operations. Evidence suggests that multisensory functions performed at subcortical levels rely less on experience compared to cortical multisensory functions (e.g., Putzar et al., 2010). As the auditory model used in our study to extract and impose statistics represented peripheral computations, from the cochlea up to the midbrain (McDermott and Simoncelli, 2011), it is possible that the functional development of the set of auditory statistics included in our stimuli is exclusive competence of the auditory system and does not rely on other modalities, such as vision. Further studies including statistics resulting from computations beyond primary cortex (Chi, Ru and Shamma, 2005; Norman-Haignere and McDermott, 2018) and involving other types of naturalistic stimuli will be able to shed light on potential visual influences over the development of higher-order auditory statistics.

### Functional interplay between selective auditory computations and vision

By comparing CB, SC, and LB groups, we can affirm that while early vision is not necessary for the typical development of the auditory statistics processing, the lack of vision can still influence the processing of local features. No compensatory effects were found in LB group for any of the tested computational modes, while a detrimental effect was present for the acoustic local features analysis.

One possibility for this mode to be affected in LB, but not CB, individuals is that a visual influence on auditory computations can only take place after the major development of the auditory system has occurred. Consistent evidence shows that, even considering basic auditory functions, human performance keeps improving gradually for over nearly a decade. Frequency discrimination, such as perceiving differences between two tones presented sequentially, does not mature until roughly 10 years of age for low frequencies (Maxon and Hochberg, 1982; Jensen and Neff, 1993; Moore et al., 2011). Similarly, the thresholds for detecting amplitude modulations become adult-like only after 10 years of age (Banai, Sabin and Wright, 2011). The use of local features comprised a number of operations, including the measurement of amplitude modulations over time (Dau, Kollmeier and Kohlrausch, 1997). Thus, it is possible they are designed to develop independently from visual input and only after their functional development is completed, interactions across senses are allowed.

The observed effects might also depend on the development of multisensory functions. Previous studies have revealed that certain aspects of multisensory integration can occur only by the age of 8-10 years (Gori et al., 2008). In the same vein, basic audio-visual (AV) multisensory facilitations have been found immature until the age of 9 years old while similar patterns have been observed between adolescents and adults (Brandwein et al., 2011). Similarly, the development of multisensory speech perception has been found to progress until late childhood (Ross et al., 2011). The present data might suggest that the interactions between basic auditory computations tested here and vision occur after full development of AV multisensory functions. The availability of visual input in early developmental phases would prompt the typical development of AV interactions. As a result, the sudden and permanent loss of visual input could, in turn, alter auditory processing which had typically developed.

Shamma (2001) suggested that the most essential auditory percepts (timbre, pitch and localization) can be derived through computational approaches typically employed to model visual processing (see also McDermott and Simoncelli., 2011). As an example, extracting the profile of sound spectrum has been associated to the type of neural computations which are necessary for the extraction of the form of an image. The common principle could rely on lateral inhibition for the enhancement of edges or peaks (Lyon and Shamma, 1996; Shamma, 1985). The present results might suggest that if a unified plan for basic auditory and visual computations exists, it might develop, for certain functions, independently for each sensory modality.

The absence of associations between the accuracy in both tasks and duration or onset of blindness (Figure S5) suggested that the performance: (i) was not influenced by the number of years people were visually deprived (in our LB sample, duration of blindness spanned from 2 years up to 28 years); (ii) was similar for people who became blind during childhood (10 years of age) as well as during adulthood (up to 51 years of age, in our LB sample). However, we had the chance to test two early blind individuals whose blindness onset occurred during their third year of life (Figure S3). Remarkably, their results systematically overlapped with CB and SC groups, providing further support to the evidence that the dampening of local features processing manifests only if sight loss occurs relatively late in the development.

Finally, the deficit observed in the LB group does not exclude that limiting local features accessibility can provide ecological advantages. Identifying a Sound Texture equals recognizing the statistics included in its sound waveform (McDermott and Simoncelli, 2011). Thus, it is possible that relying on statical averaging only when information is consistent -at the expense of temporal details-prompts sound-object recognition in an everyday environment. The results in our LB group could suggest a form of adaptive perceptual learning (Watanabe and Sasaki, 2015) relying on implementing strategical changes aimed at facing a remarkable loss in the overall available sensory input (de Villers-Sidani and Merzenich, 2011).

## CONCLUSION

Overall, this evidence has several important implications. First, it discloses how basic auditory computations can develop independently from early visual input. Second, it shows a selective detrimental effect induced by a late-onset sight loss over selective, non-spatial aspects of the auditory processing. Third, it has the potential to expand our approaches in several fields, including auditory, visual, multisensory, and brain disorders research. Moreover, it provides novel ground for applied science such as sensory substitution device and auditory rehabilitation strategies for people who lost vision compared to those born blind. Finally, we proved that combining computational methods with human models of sensory deprivation provides the context to assess the degree of plasticity of specific computations performed by the sensory systems. While the present study focused on the auditory domain, similar approaches could be employed for other modalities. Providing the resemblance of the Auditory Texture Model (McDermott and Simoncelli, 2011) with a previously validated Visual Texture Model (Portilla and Simoncelli, 2000), and the presence of textures in other modalities, such as touch (Picard et al., 2003; Weber et al., 2013), it can be possible to address with similar approaches the development and sensory interplays across computations performed by other senses.

## MATERIALS AND METHODS

Experimental procedures and methods are inspired by the work of McDermott, Schemitsch, and Simoncelli (2013).

### Synthetic texture synthesis

Synthetic textures were synthesized from the model described in a previously published paper (McDermott and Simoncelli, 2011). For synthesis, we used the MATLAB-based Sound Texture Synthesis Toolbox, available here: http://mcdermottlab.mit.edu.

Auditory statistics were computed from different signals of 7-s length, each one being an Original Recording of a natural Sound Texture, with a sampling rate of 20kHz. These sounds are also available on the website previously cited. The Original Recordings we used for synthesis and experiment were a subset of the ones used by McDermott, Schemitsch, and Simoncelli (2013). For each original recording, cochlear envelopes and modulation bands were extracted from the signal, by filtering it through the Auditory Texture Model (McDermott and Simoncelli, 2011). Precisely, extracted statistics included marginal moments (mean, variance, and skew) of each cochlear envelope, envelope cross-correlation, the normalize power in each modulation band, and two correlations between modulation bands (C1 and C2). Kurtosis was omitted from the computation (see toolbox-authors’ suggestion). Statistics were measured with a temporal weighting function that faded to zero over the first and last second of the original recordings to avoid boundary artifacts (McDermott and Simoncelli, 2011).

After setting the parameters, the process was initialized from a 5-s white noise sample. The noise signal was again filtered through the model and the original sound statistics were imposed on its cochlear envelopes, with circular boundary conditions. The resulting envelopes were multiplied by their associated fine structure and recombined into the signal. The procedure was repeated in an iterative way, until the imposed statistics measured from the synthetic signal reached a desirable signal-to-noise-ratio (SNR) of 30-dB, which represents the ratio of the squared error of a statistics class, summed across all statistics in the class, to the sum of squared statistics values of that class. The average of each statistics class was at least 20-dB (McDermott and Simoncelli, 2011). The resulting Synthetic Sound is a 5-s random signal constraint only by the over-imposed statistics of interest (Figure 1A). The procedure was repeated four times on four different white noise samples to obtain four different synthetic exemplars for each Sound Texture (adapted from McDermott, Schemitsch, and Simoncelli, 2013).

### Stimuli

Each synthetic exemplar was cut into excerpts of different durations, equally spaced on a logarithmic scale (40, 91, 209, 478, 1093, and 2500ms). A 10-ms Hanning window was applied to the beginning and the ending segments of each excerpt, to smooth signal onset and offset. All excerpts were equalized to the same root-mean-squared level (rms = 0.1).

### Experimental Procedures

Participants sat in front of a computer and performed the task using the mouse. Experiments were implemented in MATLAB. All subjects were blindfolded by a mask that could filter almost 100% of the light and kept their eyes closed. Stimuli were played on a Macbook Pro 2017, with a built-in sound card with a frequency sampling rate of 44.1 Hz, through a headphone set Audio Technica Pro ATH-M50X, at a volume of c.a. 75 dB SPL, which was kept constant for each stimulus.

The task was very similar to the one in the original paper by McDermott, Schemitsch, and Simoncelli (2013) but modified to be suitable for visually deprived individuals. In fact, no visual interaction with the screen was required and audio instructions were provided through the headphones in Italian or English, according to participant’s preferred language.

Participants performed in two sessions, each comprising either version of the two experiments (Exemplar Discrimination or Texture Discrimination), presented in a counterbalanced order across subjects. Both sessions were performed in the same day with a half-an-hour break in between; each session lasted approximately 35 -40 minutes. 54 Sound Textures were employed in Texture Discrimination and 36 in Exemplar (see Table S1, Supplementary information); each experiment comprised 216 trials. Every 54 trials, corresponding to about 6-7 minutes of stimulation, participants were allowed to take a few-minutes break, for a total of four runs per each session, separated by four small breaks.

In both sessions, each trial consisted of a triplet of sounds of the same duration. Stimuli duration could vary across trials, and six durations were employed (either 40, 91, 209, 478, 1093 or 2500ms). The number of trials per duration was equal across all the six possible durations (36 trials for each one) and stimuli were presented in a randomized order. To control for a stimulus expectancy effect, inter-stimulus interval (ISI) could vary between 400, 500, 600, 700, and 800-ms. Participants were asked to be as accurate as possible in their choice and there was no time-limit to answer. Once the participant had responded, the presentation of the next triplet occurred after a short pause lasting between 2 and 2.5-s, with steps of 100ms.

Test sessions were performed before both of the actual experimental sessions, to make sure participants understood the tasks and to have them familiarize with the type of stimuli. Test stimuli were drawn randomly across the 36 texture pairs (Table S1, Supplementary information); excerpts used in the trial session were then excluded by the actual experiment. Three trials for each duration were presented in a random order during test sessions (for a total of 18 trials per duration). A feedback was provided only during the trial session and consisted in an audio message stating if their response was correct or incorrect. Feedback was not provided during experimental sessions, following the protocol by McDermott, Schemitsch, and Simoncelli, 2013.

### Exemplar Discrimination

Two excerpts coming respectively from two exemplars of the same Sound Texture were selected. Both excerpts were extracted at the same time point along the 5-s segments. Excerpts were then presented as triplet of sounds in a trial: consequently, one of the two excerpts was repeated twice, while the third sound was the other. The different sound could be located as the first one or the last one in the triplet, and this varied randomly across trials.

Participants were informed that two stimuli will be identical and were asked to indicate which stimulus was different from the other two. If they thought it was the first one, they would click the left button of the mouse, otherwise the right one. The middle sound was used as the reference (Figure 1C).

### Texture Discrimination

Three excerpts were drawn among three different exemplars coming from pairs of Original Sound Textures: two of the excerpts came from two different exemplars of the same Sound Texture while the third one from an exemplar of another Sound Texture. Sound Textures were paired according to similarity, as done by McDermott, Schemitsch, and Simoncelli, 2013; Table S1, Supplementary information) and the different exemplar was a synthetic sound derived from the same white noise of one of the other two exemplar, so their original associated fine structure was the same, while only the imposed statistics were different. All three excerpts were extracted at the same time point along the 5-s segments. Again, the three excerpts were presented in a trial so that all the sounds were different, but two of them would originate from the same Sound Texture, while the other one had a different sound source. Participants were made aware of the fact that all sounds could be different and were asked to report which one came from the diverging acoustic source. Some examples were provided in order to facilitate task comprehension (i.e., “Two sounds can be the sound of a fireplace, the other one is the sound of the rain”). The correct answer could be either the first or last sound in the triplet, while middle one had to be used has a reference. Middle excerpt and deviant excerpts were derived from the same white noise sample. If participants thought the deviant was the first sound, they would click the left button of the mouse, otherwise the right one (Figure 1B).

### Average statistical variability

Standard deviation of a set of employed envelope statistics (envelope mean, variance, skew, cross-correlation, and modulation power) was measured between couples of excerpts presented in both experiments. The pair could be made of excerpts coming from different exemplar of the same Sound Texture (column 1, Table S1, Supplementary information) or excerpts coming from different Sound Textures (column 1 and 2, Table S1, Supplementary information; Figure 1D). The average of all statistics variability among couples of sounds was computed to make predictions about performance. When both excerpts originated from different exemplars of the same Sound Textures, their variability tended toward zero as duration increased, whereas opposite trend was observed when stimuli came from different Sound Textures. Moreover, differences in variability between the two conditions (Different Exemplar vs. Different Texture) increased with duration (Figure 1D). Separately for each of the aforementioned statistics, we performed pairwise comparisons (t-tests, FDR corrected) between the two conditions (Different Exemplar vs. Different Texture) and observed that variability between couples of excerpts coming from different Sound Textures, as compared to ones originating from different exemplar of the same Texture in envelope mean was larger for all durations (all p > 0.05, corrected); in envelope skewness in was larger at duration 91, 478, 1093, 2500ms (all p > 0.05, corrected); variability significantly diverged only at long durations in envelope variance, envelope cross-correlation (478ms, 1093, and 2500ms; all p < 0.001, corrected), and modulation power (1093, 2500ms; all p < 0.01, corrected; Figure S1). Analyses were performed using MATLAB.

### Participants

Data from three groups of participants, matched for size, gender, and age, were analyzed in this study. We recruited a group of congenitally blind individuals (CB; N = 18; F = 9; mean age = 37.06 years; std = 10.75). Data of the 18 CB individuals were used as reference for the recruitment of a group of late blind individuals and a group of sighted controls. Each CB individual in our sample was matched with a sighted individual and a LB individual of same gender and similar age (mean age of groups was within 2 std of difference), for a total of 18 participants for each group. Late onset blind individuals (LB; N =18; F = 9; mean age = 40.11; std = 12.68); range of blindness onset= 10-51 years; range of blindness duration= 2-28 years), and sighted controls (SC; N = 18; F = 9; mean age = 38.06 years; std = 12.93).

Before experimental session began, all blind participants underwent a short interview to gather several information, especially about onset, cause, and duration of blindness, together with other anamnestic information.

All participants in the final sample were healthy and fully understood the task requests. During recruitment, exclusion criteria comprised documented hearing impairment (i.e., acoustic implants, tinnitus), neurological disorders. The data of 3 CB individuals, 1 early blind participant (blindness onset: 7 months) and of 2 SC were not included in the final sample and thus were not analyzed due to their inability to perform/terminate one or both sessions or being easily distracted during experimental sessions (i.e., participant often asked questions and talked during the task). For all of the blind participants in the sample, blindness was total and caused only by peripherical pathologies (Table 1).

Two early blind (EB) participants were also tested (EB1, gender = M; age = 26, blindness onset = 3years; EB2, gender = M; age = 24, blindness onset= 3 years). Results from these participants were excluded from the analyses, as blindness onset was borderline between blind groups. Their data are plotted in Supplementary information (Figure S3).

Beforehand, all participants were informed about the procedures and purpose of the study and signed a written informed consent prior to testing. The study was approved by the regional Ethical Committee (CEAVNO protocol n 24579). The study protocol adhered to the guidelines of the Declaration of Helsinki (2013).

### Statistical Analyses

Proportion of correct responses for each individual was used as dependent measure for statistical analyses. To assess that sample data were normally distributed, we performed both Kolmogorov-Smirnov and Shapiro-Wilk tests with the average of proportion of correct answers in each experiment (separately) as dependent variables. In each test, dependent variable was split according to group. Both tests indicated that data were normally distributed in both experiments and for all of the groups, giving significance values higher than 0.05 (see Supplementary information, Table S2).

We performed an ANCOVA using IBM SPSS Statistics for Macintosh, Version 26.0. The model included the between-participants factor Group (SC, CB, LB) and two within-participant factors: Experiment (Exemplar Discrimination vs. Texture Discrimination) and Duration (40, 91, 209, 478, 1093, 2500); given the relatively large age-range across the entire sample (20-62 years old), age was included as a covariate. Since groups were matched, age unlikely accounted for between-groups effects, but it could still have had an impact at the individual level.

Mauchly’s test indicated that the assumptions of sphericity had been violated for the interaction Experiment * Duration (c2(2) = 25.62, p < 0.05). Thus, degrees of freedom were corrected using Greenhouse-Geisser estimates of sphericity (ε = 0.81).

There was a significant main effects of Duration, F_(5, 250)_ = 9.45, p < 0.001, η^2^ = 0.16, and its interaction with the factor Experiment, F_(4.07, 203.35)_ = 15.91, p < 0.001, η^2^ = 0.24. Also, there was a significant main effect of the between-participants factor Group, F_(2, 50)_ = 4.99, p = 0.01, η^2^ = 0.17, and a significant interaction between factors Group and Experiment, F_(2,50)_ = 8.35, p = 0.001, with a large effect size of η^2^ = 0.25. Moreover, there was a significant interaction effect among Experiment, Duration, and Group, F_(10, 250)_ = 3.29, p = 0.001, with a medium-to-large eta-squaredof η^2^ = 0.12. Participant’s age and its interactions with all other independent variables in the model were non-significant (all p > 0.14).

To break down the three-way interaction, we performed planned pairwise comparisons, using two-sided t-tests only on pre-specified effects of interest, as opposed to investigating all main effects and interactions as in exploratory analyses (see Cramer et al., 2016).

The pre-specified contrasts were: (i) Within Group contrasts, for each group (CB, LB, SC) and duration (40, 91, 209, 478, 1093, 2500): Exemplar Discrimination vs. Texture Discrimination), for a total of 18 contrasts (Figure 2A, 2B, 2C). (ii) Between Group contrasts (within durations and experiments). For each duration (40, 91, 209, 478, 1093, 2500) and experiment (Exemplar Discrimination vs. Texture Discrimination): LB vs. SC, CB vs. LB, LB vs. CB, for a total of 36 contrasts (Figure 2D, 2E). (iii) Within Experiment contrasts (within Group and across duration). For each experiment (Exemplar Discrimination and Texture Discrimination) and group (CB, LB, SC): 40 vs. either 91, 209, 478,1093 or 2500; 91 vs. either 209, 478,1093 or 2500; 209 vs. either 478,1093 or 2500; 478 vs. either 1093 or 2500; 1093 vs. 2500. For a total of 90 contrasts. P-values were corrected for multiple comparisons across all 144 pre-specified pairwise comparisons (the contrasts of interest listed above) using the false discovery rate (FDR; Benjamini, Drai, Elmer, Kafkafi, and Golani, 2001) and a q-value of 0.05. Post-hoc comparisons and adjusted p-values were computed using RStudio version 1.2.1335.

### Within Group contrasts

#### Sighted controls

For the SC group, the within Group contrasts Exemplar Discrimination vs. Texture Discrimination were significantly different at durations 40 (p < 0.001, corrected), 91 (p < 0.01, corrected), 478 (p < 0.05, corrected), 1093 (p < 0.001, corrected), and 2500 (p < 0.001, corrected). For short durations (40, 91), performance was higher in Exemplar Discrimination (mean proportion correct: 40ms = 0.87; SE = 0.02; 91ms = 0.89; SE = 0.02) compared to Texture Discrimination (mean proportion correct: 40ms = 0.65; SE= 0.02; 91ms = 0.77; SE = 0.02). Conversely, for longer durations (478, 1093, 2500), performance was better for Texture Discrimination (mean proportion correct: 478ms = 0.91; SE = 0.01; 1093ms, 0.97; SE = 0.01; 2500ms = 0.97; SE = 0.01) compared to Exemplar Discrimination (mean proportion correct: 478ms = 0.84; SE = 0.02; 1093ms, 0.77; SE = 0.03; 2500ms = 0.70; SE = 0.03). No Difference was observed at duration 209 (mean proport
ion correct at 209: Exemplar = 0.88; SE = 0.02; Texture = 0.85; SE = 0.01; p > 0.05, corrected; Figure 2A).

### Congenitally blind group

For the CB group, results pattern was remarkably similar to SC’s. The contrasts Exemplar Discrimination vs. Texture Discrimination were significant at duration = 40 (p < 0.001, corrected), 91 (p < 0.01, corrected), 478 (p < 0.001, corrected), 1093 (p < 0.001, corrected), and 2500 (p < 0.001, corrected), with performance being higher for short durations in Exemplar Discrimination (mean proportion correct: 40ms = 0.87; SE = 0.02; 91ms = 0.85; SE =0.03) compared to Texture Discrimination (mean proportion correct: 40ms = 0.65; SE = 0.02; 91ms = 0.77; SE = 0.02). On the contrary, for long durations, performance was higher in Texture Discrimination (mean proportion correct: 478ms = 0.92; SE = 0.01; 1093ms, 0.97; SE= 0.01; 2500ms = 0.98; SE = 0.01) compared to Exemplar Discrimination (mean proportion correct: 478ms = 0.83; SE= 0.02; 1093ms = 0.77; SE = 0.03; 2500ms = 0.64; SE= 0.04). As for the SC group, no Difference was observed at duration = 209 (mean proportion correct at 209: Exemplar Discrimination = 0.87; SE = 0.02; Texture Discrimination = 0.86; SE = 0.02; p > 0.05, corrected; Figure 2B).

### Late-onset blind group

For the LB group, comparisons between Exemplar Discrimination vs. Texture Discrimination revealed a substantially different pattern of results compared to CB and SC groups. The performance of the two tasks did not differ at short durations 40 (mean proportion correct: Exemplar Discrimination = 0.70; SE = 0.03; Texture Discrimination = 0.69; SE= 0.01), 91 (mean proportion correct: Exemplar Discrimination = 0.76; SE = 0.03; Texture Discrimination = 0.74; SE = 0.02), 209 (mean proportion correct: Exemplar Discrimination = 0.85; SE = 0.02; Texture Discrimination = 0.79; SE = 0.01; all p > 0.07, corrected). Significant differences emerged only at longer durations 478 (p < 0.001, corrected), 1093 (p < 0.001, corrected), and 2500 (p < 0.001, corrected), where performance was better for Texture Discrimination (mean proportion correct: 478ms = 0.91; SE= 0.01; 1093ms = 0.97; SE = 0.04; 2500ms = 0.99; SE = 0.02) compared to Exemplar Discrimination (mean proportion correct: 478ms = 0.73; SE = 0.01; 1093ms = 0.72; SE = 0.03; 2500ms = 0.61; SE = 0.02; Figure 2C).

### Between Group contrasts

Contrasting SC vs. CB, no significant difference in the proportion of correct answers was observed in either Experiments (Exemplar Discrimination and Texture Discrimination) and for any of the durations (all p > 0.37, corrected).

In the Exemplar Discrimination, LB vs. SC and LB vs. CB contrasts were significantly different at specific durations: comparisons between LB and SC groups were significant at duration = 40 (p < 0.001, corrected), 91 (p < 0.001, corrected), 209 (p < 0.05, corrected), 478 (p < 0.05, corrected), and 2500 (p < 0.05, corrected); comparisons between LB vs. CB groups were significantly different at duration 40 (p < 0.001, corrected), 91 (p < 0.05, corrected), 209 (p < 0.05, corrected), and 478 (p < 0.05, corrected; Figure 2D).

In the Texture Discrimination experiment, LB vs. SC contrasts were not significantly different (all p > 0.07, corrected). A significant contrast was observed at duration 40 when comparing LB vs. CB (p < 0.05, corrected) with LB performing better than CB. None of the other contrasts at the remaining durations was significant (all p > 0.05, corrected; Figure 2E).

### Within Experiment contrasts

Overall, in the Exemplar Discrimination, performance tended to decrease with duration as in McDermott, Schemitsch, and Simoncelli (2013). for both SC and CB groups, but not for the LB group. For the SC group, the following comparisons between durations were significantly different: 1093 vs. 40, 1093 vs. 91, 1093 vs. 209, 1093 vs. 478 and 2500 vs. 40, 2500 vs. 91, 2500 vs. 209, 2500 vs. 478 (all p < 0.05, corrected) whereas the other comparisons 40 vs 91, 40 vs 209, 40 vs. 478, 91 vs. 209, 91 vs. 478, 209 vs. 478 were not significant (all p > 0.05, corrected). Similarly, for the CB group, the following comparisons resulted significant: 40 vs. 1093, 40 vs. 2500, 91 vs. 2500, 209 vs. 1093, 209 vs. 2500ms, 478 vs. 2500, and 1093ms vs. 2500ms (all p < 0.05, corrected). For the LB group, comparisons between durations were all non-significant (all p > 0.91, corrected), apart from the comparisons between duration 2500 and all of the others (40 vs. 2500, 91 vs. 2500, 209 vs. 2500, 478 vs. 2500, 1093 vs. 2500) which were significantly different (all p < 0.01, corrected).

The data for Texture Discrimination replicated the one from McDermott, Schemitsch, and Simoncelli (2013) for all groups, with performance progressively increasing with duration. For all the three groups, the following comparisons across durations were significantly different: 40 vs. either 91, 209, 478,1093, or 2500; 91 vs. either 209, 478,1093 or 2500; 209 vs. either 478,1093 or 2500; 478 vs. either 1093 or 2500 (SC: all p < 0.01, corrected; CB: all p < 0.02, corrected; LB: all p < 0.02, corrected), while the comparison between the two longest durations, 1093 vs. 2500, was significant only for the LB group and not for SC and CB groups (LB: p = 0.007, corrected; SC: p = 1, corrected; CB: p = 0.36, corrected).

### Difference in performance between Exemplar and Texture Discrimination

For every duration, each participant’s accuracy-score in Texture discrimination was subtracted from the scores in Exemplar Discrimination (Figure 2F). To test whether there was a significant difference among groups, we ran an ANOVA for repeated measures with one within-subjects factor Duration and one between-subjects factor Group. The dependent variable was the relative difference between the accuracy scores in Exemplar Discrimination and Texture Discrimination. Main effects of Group, F_(2, 51)_ = 8.72, p < 0.001, η^2^ = 0.26, and Duration, F_(5, 255)_ = 230.42, p < 0.001, η^2^ = 0.82, were significant, together with their interaction, F_(10, 255)_ = 3.22, p < 0.001, η^2^ = 0.11. We ran FDR corrected pairwise comparisons (two-tailed t-tests; q-value = 0.05) on pre-selected contrasts of interest highlighting the differences between groups within each duration. All of the comparisons between CB and SC were not significant (all p > 0.32, corrected). Comparisons between LB and CB were significant at duration 40 (p < 0.001, corrected), 478 (p < 0.03 corrected) and not significant for other durations (all p > 0.09, corrected). Finally, comparisons between SC and LB were significant for most of the durations (40, 91, 209, 478, 2500; all p < 0.03, corrected) but 1093 (p = 0.32, corrected) (Figure 2D). Absolute values of the average across participants of the differences at the Group level are displayed separately for each group in Supplementary information, Figure S4(A, B, C). For both SC (Figure S4A) and CB (Figure S4B) we observed a U-shaped trend, consisting of higher values at short and long durations and almost zero at intermediate one. On the other hand, in LB groups (Figure S4C) a different trend was observed, with values being almost zero at short durations (40, 91) and progressively increasing at longer ones.

### Testing for Disposition bias

As participants were asked to choose between first and third intervals, the temporal connotation of this 2AFC protocol could have led to a response disposition bias (e.g., listeners could have shown a tendency toward reporting one of the two intervals, for example the last one). In order to rule out that systematic trends for bias could differ across groups and, to some extent, could account for the impaired performance in LB, for each participant we calculated how many times they overestimated stimuli in one position, as compared to the other. Separately for each experiment and for each duration, the total number of times participants pressed the mouse’s left button, stating that deviant sound was the first interval, was subtracted from the total number of times that correct answer was actually the left one. Positive numbers would indicate an overestimation of stimuli in the last interval, whereas negative values would refer to an overestimation of first intervals. In order to check for significant differences, we ran a Repeated-Measure ANOVA with total number of overestimated stimuli in one position as dependent variable, Group as between-subjects factor, and Experiment and Duration as within-subjects factors. As Mauchly’s test indicated that the assumptions of sphericity had been violated for the effect Duration (c2(2) = 76.28, p < 0.001), the interactions Experiment*Duration (c2(2) = 78.76, p < 0.001), degrees of freedom were corrected using Greenhouse-Geisser estimates of sphericity (Duration, ε = 0.58; Experiment*Duration, ε = 0.56). No significant effects were found for any within-subjects factors, nor for the between-subject factor Group and their interactions (all F > 0.07). Statistics were carried out using IBM SPSS Statistics for Macintosh, Version 26.0. Data are displayed in Figure S2A.

### Testing for Learning Effect

Sessions were divided into four runs of 54 trials each. In order to check for the occurrence of divergent learning and/or tiredness effects between groups, within each experiment and duration, total number of correct answers in run 1 was used as a baseline and was subtracted from the number of correct answers in the next runs (run 2, 3, and 4). Positive values meant that performance increased compared to the first run, showing a learning effect, whereas negative values were associated with a decrease in the performance, possibly a tiredness effect. We ran a Repeated-Measure ANOVA using IBM SPSS Statistics for Macintosh, Version 26.0, with baseline-subtracted correct answers as the dependent variable, Group as between-subjects factor, and Experiment, Duration, and Run within-subjects factors. Mauchly’s test indicated that the assumptions of sphericity had been violated for the main effect of Duration (c2(2) = 67.35, p < 0.001), the interactions Experiment*Duration (c2(2) = 38.80, p < 0.001), Duration*Run (c2(2) = 150.31, p < 0.001), and Experiment*Duration*Run (c2(2) = 212.37, p < 0.001). Thus, degrees of freedom were corrected using Greenhouse-Geisser estimates of sphericity (Duration, ε = 0.68; Experiment*Duration, ε = 0.77; Duration*Run, ε = 0.62; Experiment*Duration*Run, ε = 0.49). We observed significant effects of Duration, F_(3.4, 173.47)_ = 20.11, p < 0.001, η^2^ = 0.28, Run, F_(2, 102)_ = 12.48, p < 0.001, η^2^ = 0.20, and the interactions Duration*Run, F_(10, 315.08)_=13.81, p < 0.001, η^2^ = 0.21, and Duration*Run*Group, F_(12.36, 315.08)_ = 20.11, p < 0.05, η^2^ = 0.08. Pairwise comparisons were carried out, to test whether between groups differences (SC vs. CB, CB vs. LB, SC vs. LB) existed for each run and duration. No significant difference was observed between the 54 contrasts of interest (all p > 0.05, corrected). Data are plotted in Figure S2B, showing similar trends across groups for all durations.

### Correlation between LB’s performance with Onset and Duration of blindness

We performed linear correlations between LB participants’ onset of blindness and duration of blindness with (1) their overall performance in each experiment (Supplementary information, Figure S5A) and (2) their performance for each duration in each experiment (Figure S5B and S5C). Pearson’s correlation coefficient (RHO) between each pair pairwise comparison was computed, together with p-values. We observed no significant correlation for most of the conditions and the variable tested (all p > 0.05). Correlations were performed and plotted with MATLAB.

## ACKNOWLEDGMENTS

We want to thank Antonio Quatraro, Istituto per la Ricerca la Formazione e la Riabilitazione (IRIFOR) in Florence, Unione Italiana Ciechi (UIC) in Lucca, Pisa, Pistoia, Prato, Trento, and Istituto Cavazza in Bologna. Finally, we are grateful to Dr. Peter Neri for providing insights on the analyses and interpretation of the results. Funding: Davide Bottari (PRIN 2017 research grant. Prot. 20177894ZH)

