## Supplementary Material for "Interactions between auditory statistics processing and visual experience emerge only in late development"

All participants in the final sample were healthy and fully understood the task requests. During recruitment, exclusion criteria comprised documented hearing impairment (i.e. acoustic implants, tinnitus), neurological disorders, attentional deficits, the inability of participant to perform/terminate one or both sessions or being easily distracted during experimental sessions (i.e. noisy environment, participant often asked questions and talked during performing the task). For all of the blind participants in the sample, blindness was total and caused only by peripherical pathologies (**Table S2**).

Two early blind (EB) participants were also tested (EB1, gender = M; age = 26, blindness onset = 3years; EB2, gender = M; age = 24, blindness onset= 3 years).

Results from these participants were excluded from the analyses, as blindness onset was borderline between blind groups. Their data are plotted in **Figure S1**.

Beforehand, all participants were informed about the procedures and purpose of the study and signed a written informed consent prior to testing. The study was approved by the regional Ethical Committee (CEAVNO protocol n 24579). The study protocol adhered to the guidelines of the Declaration of Helsinki (2013).

### QUANTIFICATION AND STATISTICAL ANALYSIS

Proportion of correct responses for each individual was used as dependent measure for statistical analyses.
We performed an ANCOVA using IBM SPSS Statistics for Macintosh, Version 26.0. The model included the between-participants factor Group (SC, CB, LB) and two within-participant factors: Experiment (Exemplar Discrimination vs. Texture Discrimination) and Duration (40, 91, 209, 478, 1093, 2500); given the relatively large age-range across the entire sample (20-62 years old), age was included as a covariate. Since groups were matched, age unlikely accounted for between-groups effects, but it could still have had an impact at the individual level.

Mauchly's test indicated that the assumptions of sphericity had been violated for the interaction Experiment * Duration ($c^2_{(2)} = 25.62$, $p = 0.03$). Thus, degrees of freedom were corrected using Greenhouse-Geisser estimates of sphericity ($e = 0.81$).
There was a significant main effects of Duration, $F_{(5, 250)} = 9.45$, $p < 0.001$, $\eta^2 = 0.16$, and its interaction with the factor Experiment, $F_{(4.07, 203.35)} = 15.91$, $p < 0.001$, $\eta^2 = 0.24$. Also, there was a significant main effect of the between-participants factor Group, $F_{(2, 50)} = 4.99$, $p = 0.01$, $\eta^2 = 0.17$, and a highly significant interaction between factors Group and Experiment, $F_{(2,50)} = 8.35$, $p = 0.001$, $\eta^2 = 0.25$. More importantly, there was a highly significant interaction effect among Experiment, Duration, and Group, $F_{(10, 250)} = 3.29$, $p = 0.001$, $\eta^2 = 0.12$. Participant's age and its interactions with all other independent variables in the model were non-significant (all $p > 0.14$).

To break down the three-way interaction, we performed planned pairwise comparisons, using t-tests only on pre-specified effects of interest, as opposed to investigating all main effects and interactions as in exploratory analyses (see Cramer et al., 2016).
The pre-specified contrasts were: (i) Within Group contrasts, for each group (CB, LB, SC) and duration (40, 91, 209, 478, 1093, 2500): Exemplar Discrimination vs. Texture Discrimination), for a total of 18 contrasts (see Figure 2A, 2B, 2C). (ii) Between Group contrasts (within durations and experiments). For each duration (40, 91, 209, 478, 1093, 2500) and experiment (Exemplar Discrimination vs. Texture Discrimination): LB vs. SC, CB vs. LB, LB vs. CB, for a total of 36 contrasts (see Figure 2D, S2E). (iii) Within Experiment contrasts (within Group and across Duration). For each experiment (Exemplar Discrimination and Texture Discrimination) and group (CB, LB, SC): 40 vs. either 91, 209, 478,1093 or 2500; 91 vs. either 209, 478,1093 or 2500; 209 vs. either 478,1093 or 2500; 478 vs. either 1093 or 2500; 1093 vs. 2500. For a total of 90 contrasts (see Figure S2A, S2B). P-values were corrected for multiple comparisons across all 144 pre-specified pairwise comparisons (the contrasts of interest listed above) using the false discovery rate (FDR; Benjamini, Drai, Elmer, Kafkafi, and Golani, 2001) and a q-value of 0.05. Post-hoc comparisons and adjusted p-values were computed using RStudio version 1.2.1335.

#### Within Group contrasts (within Duration and across Experiment)

##### *Sighted controls*

For the SC group, the within Group contrasts Exemplar Discrimination vs. Texture Discrimination were significantly different at durations 40 ($p < 0.001$, corrected), 91 ($p <$

On the other hand, in LB groups (**Figure S2C**) a different trend was observed, with values being almost zero at short durations (40, 91) and progressively increasing at longer ones.

**Average statistical variability**

We ran a Repeated-Measure ANOVA using IBM SPSS Statistics for Macintosh, Version 26.0, with baseline-subtracted correct answers as dependent variable, Group as between-subjects factor and Experiment, Duration, and Run within-subjects factors. Mauchly's test indicated that the assumptions of sphericity had been violated for the main effect of Duration ($c^2_{(2)} = 67.35$, $p < 0.001$), the interactions Experiment*Duration ($c^2_{(2)} = 38.80$, $p < 0.001$), Duration*Run ($c^2_{(2)} = 150.31$, $p < 0.001$), and Experiment*Duration*Run ($c^2_{(2)} = 212.37$, $p < 0.001$). Thus, degrees of freedom were corrected using Greenhouse-Geisser estimates of sphericity. We observed significant effects of Duration, $F_{(3.4,\ 173.47)} = 20.11$, $p < 0.001$, $\eta^2 = 0.28$, Run, $F_{(2,\ 102)} = 12.48$, $p < 0.001$, $\eta^2 = 0.20$, and the interactions Duration*Run, $F_{(10,\ 315.08)} = 13.81$, $p < 0.001$, $\eta^2 = 0.21$, and Duration*Run*Group, $F_{(12.36,\ 315.08)} = 20.11$, $p < 0.05$, $\eta^2 = 0.08$. Pairwise comparisons were carried out, in order to test whether between groups differences (SC vs. CB, CB vs. LB, SC vs. LB) existed for each run and duration [3]. No significant difference was observed between the 54 contrasts of interest (all $p > 0.05$, corrected). Data are plotted in **Figure S4B**, showing similar trends across groups for all durations.

**Correlation between LB's performance with Onset and Duration of blindness**

We performed linear correlations between LB participants' onset of blindness and duration of blindness with (1) their overall performance in each experiment (**Figure S5A**) and (2) their performance for each duration in each experiment (**Figure S5B and S5C**).

Pearson's correlation coefficient (RHO) between each pair pairwise comparison was computed, together with p-values. We observed no significant correlation for most of the conditions and the variable tested (all $p > 0.05$).

Correlations were performed and plotted with MATLAB.

### RESOURCE AVAILABILITY

**Lead Contact**

**Data and code availability**

Synthesis algorithm is available at: http://mcdermottlab.mit.edu/downloads.html

Analysis scripts by contacting the authors.

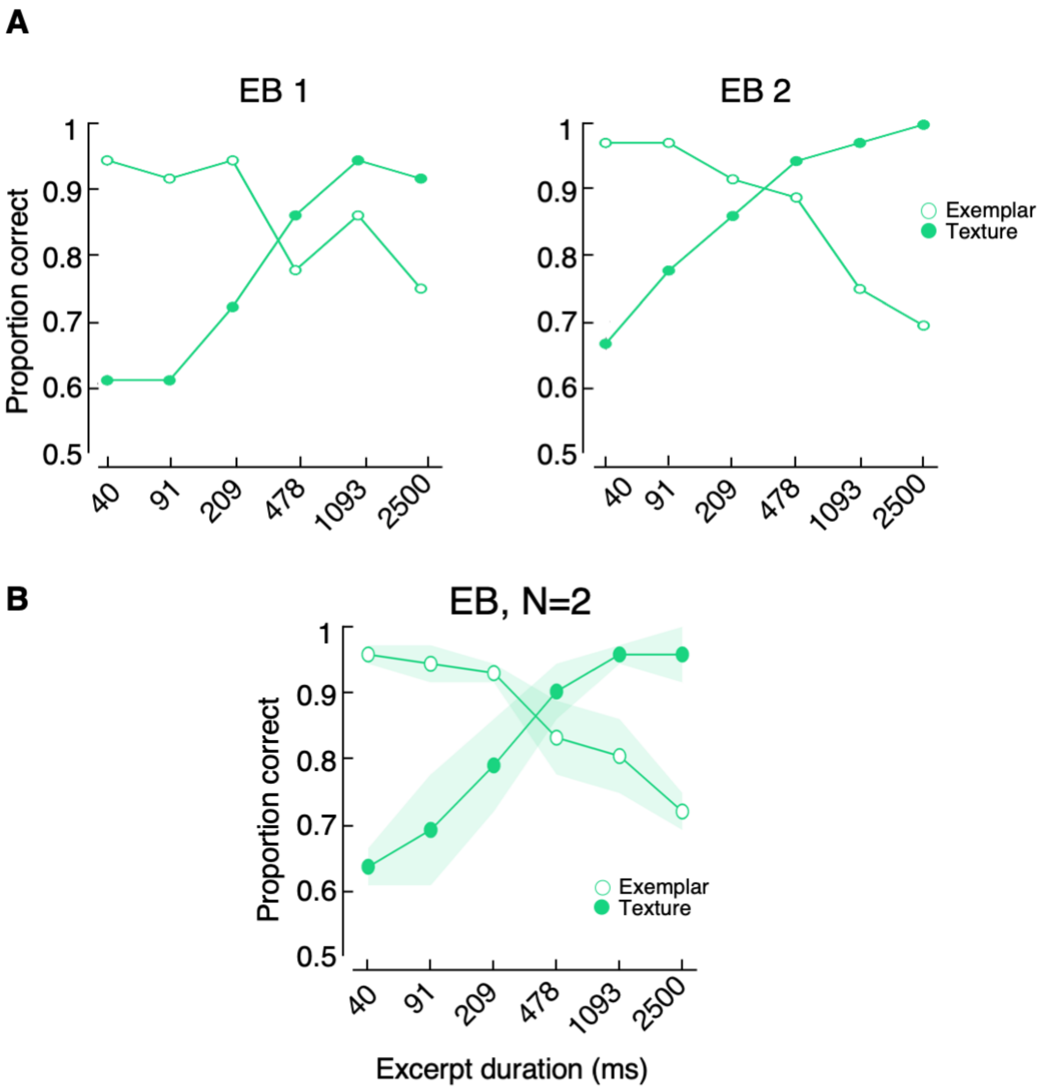


**Figure S1. Early Blinds performance.**

A) Two early blinds participants were tested (onset of blindness < 3yo). For each EB participants proportion of correct answers is plotted separately for each experiment.

B) Mean proportion of correct across both EB participants. Results are comparable to the ones of SC and CB groups. Shaded regions indicate interpolated standard error of the mean (SE) at each mean point.

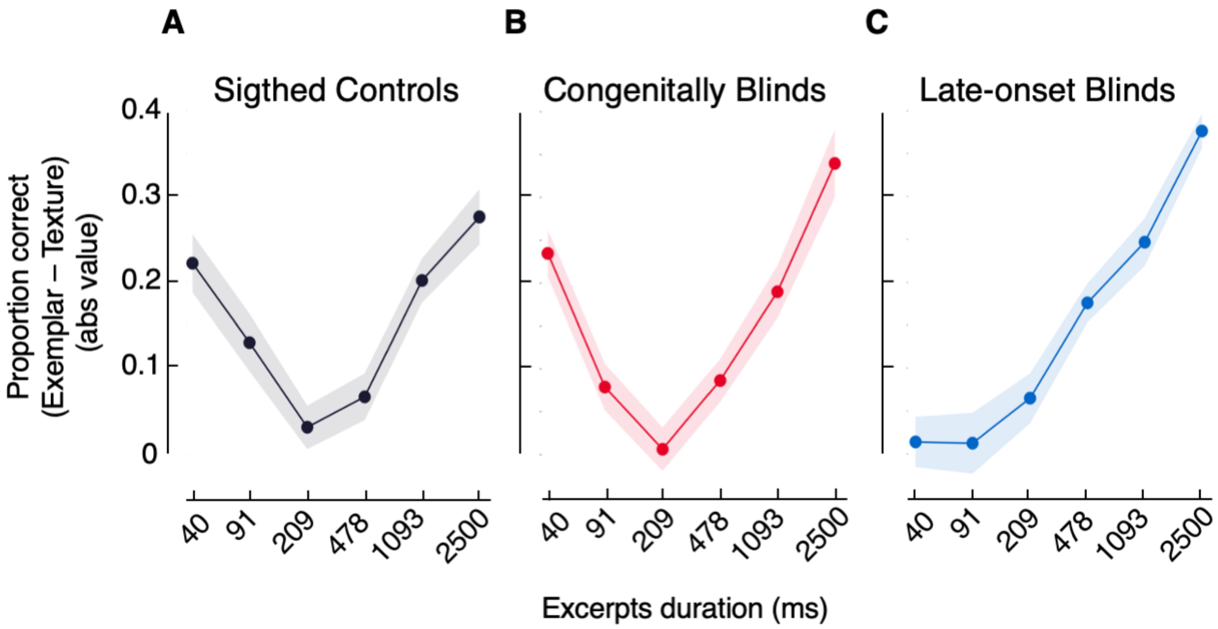


**Figure S2. Mean Difference between Experiments**

A, B, C) Absolute values of the average difference between proportion of correct answers in Exemplar Discrimination vs. Texture discrimination at the Group level is displayed separately for each duration for SC (A), CB (B), and LB (C). Both SC and CB groups show a fundamentally similar U-shaped trend, with higher difference for short and long durations and almost no difference for intermediate ones, while LB group display a different trend at short durations where there was no difference in the way the two tasks were performed. Shaded areas show interpolated standard error of the mean (SE) at each mean point.

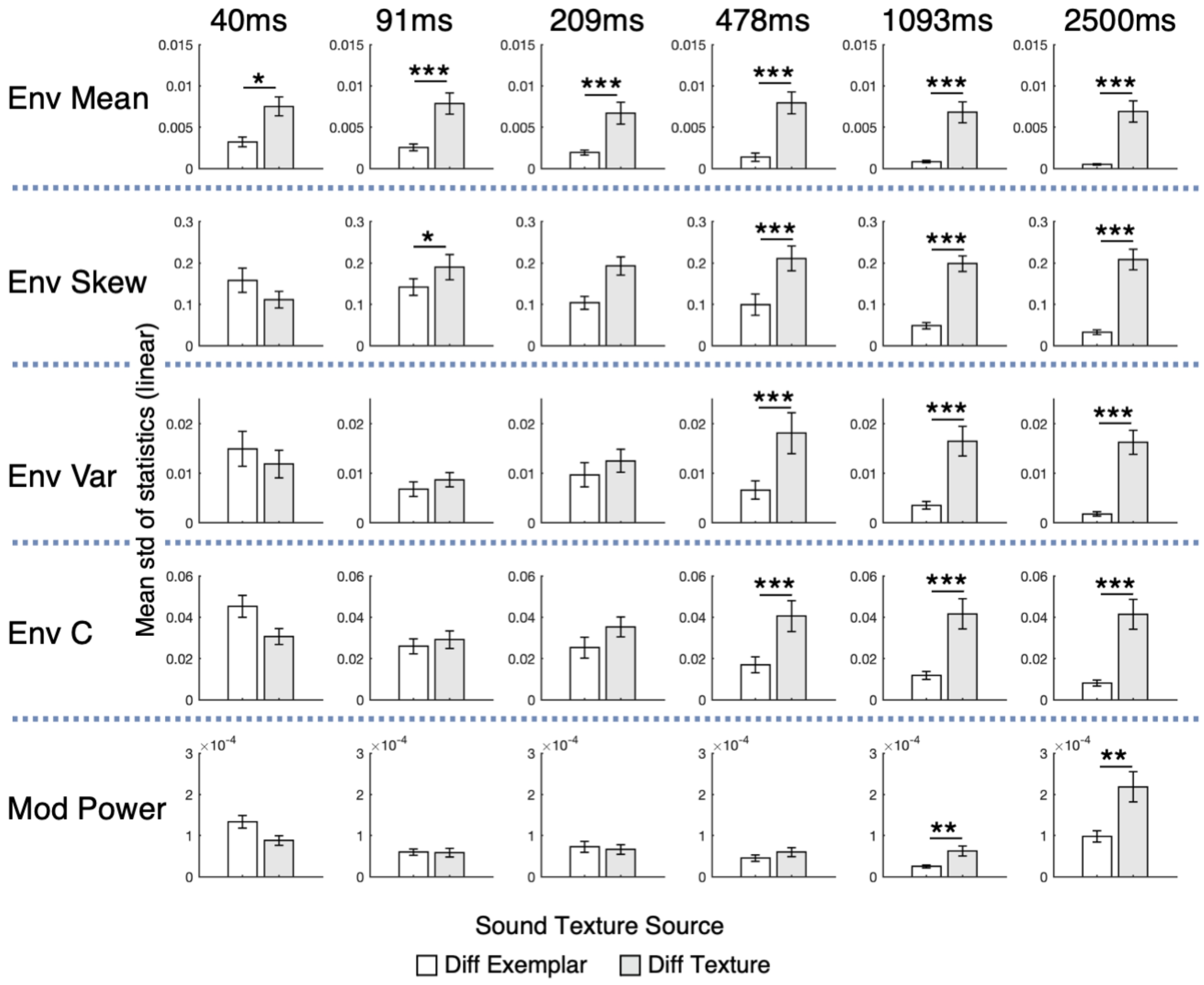


**Figure S3. Statistical variability**

For every statistic separately, we performed pairwise t-tests comparisons between standard deviation of couples originating from two different exemplars of the Same Texture as compared to excerpts originating from different Sound Textures. For majority of statistics, significant differences were observed specifically at long durations, accounting for results in all three groups in all three groups in Texture Discrimination and validating statistical averaging theory. Results were corrected for FDR. *** $p < 0.001$; ** $p < 0.01$; * $p < 0.05$.

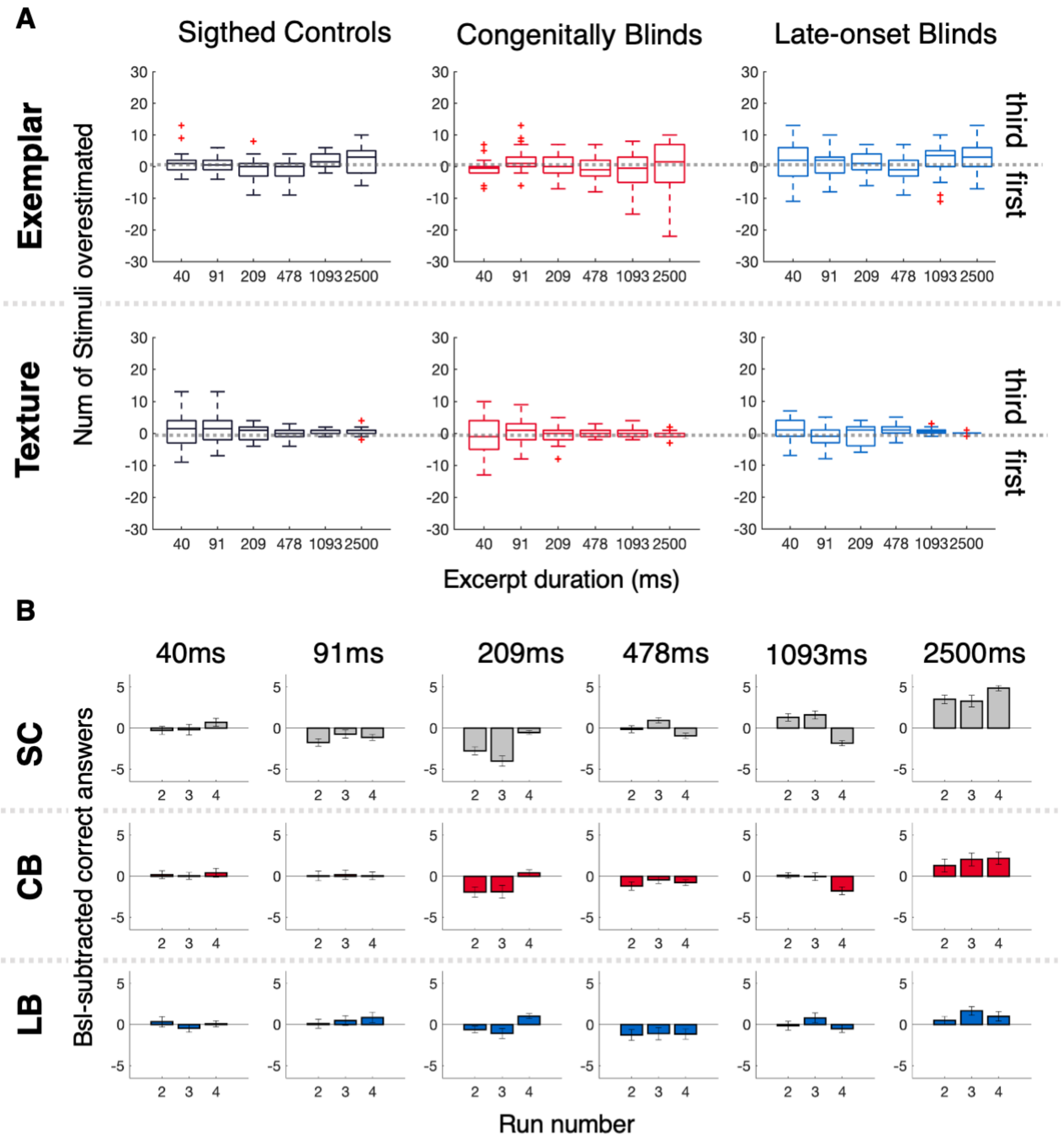


**Figure S4. Disposition bias and Learning effects do not differ between Groups.**
A) Disposition bias. For each participant, the total number of times they responded “first” was subtracted from the total number of times the deviant stimulus was actually in the first position. Median and interquartile range are plotted for each Group, Experiment, and Duration. Positive numbers represent times in which listeners overestimated the third sound; negative numbers, the first.
For each experiment and duration there was no significant difference across groups.
B) Learning effect. For each participant, the number of correct answers in run 1 (baseline) were subtracted from the correct answers of the other three runs (2, 3, and 4), each run consisting of 54 trials. Mean difference for run 2, 3, and 4 across groups and for each duration are displayed.
Errorbars represent standard error of the mean (SE) at each point.

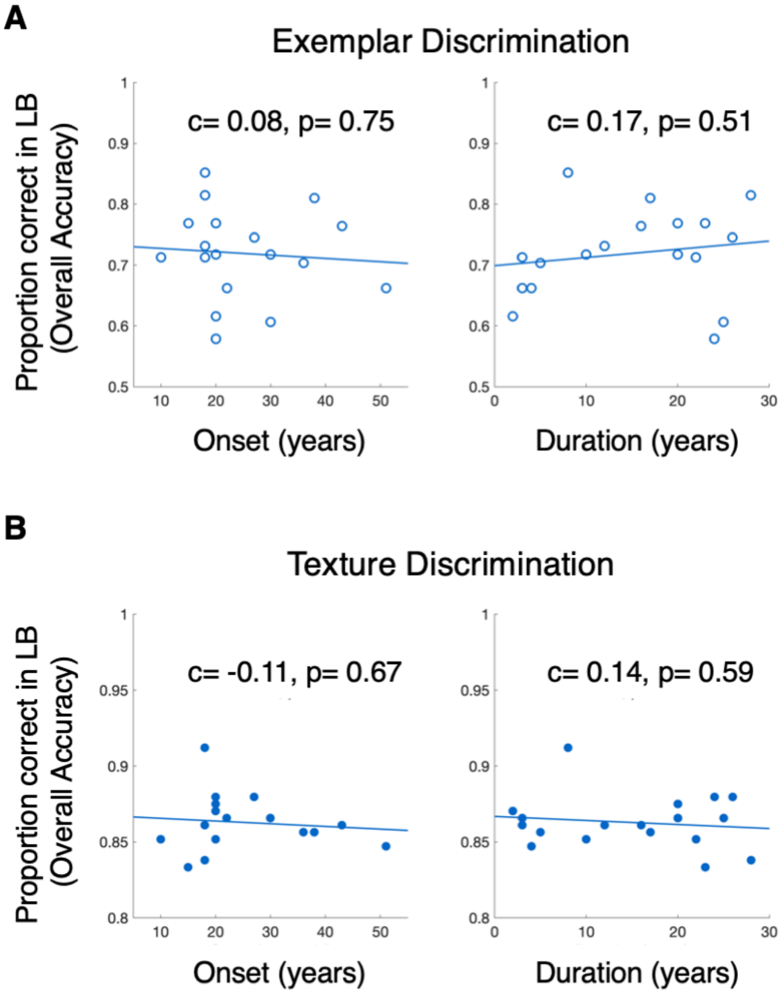
A
Exemplar Discrimination
c= 0.08, p= 0.75
c= 0.17, p= 0.51
Proportion correct in LB (Overall Accuracy)
Onset (years)
Duration (years)
B
Texture Discrimination
c= -0.11, p= 0.67
c= 0.14, p= 0.59
Proportion correct in LB (Overall Accuracy)
Onset (years)
Duration (years)

**c**

Correlation Performance Exemplar with Onset and Duration of Blindness

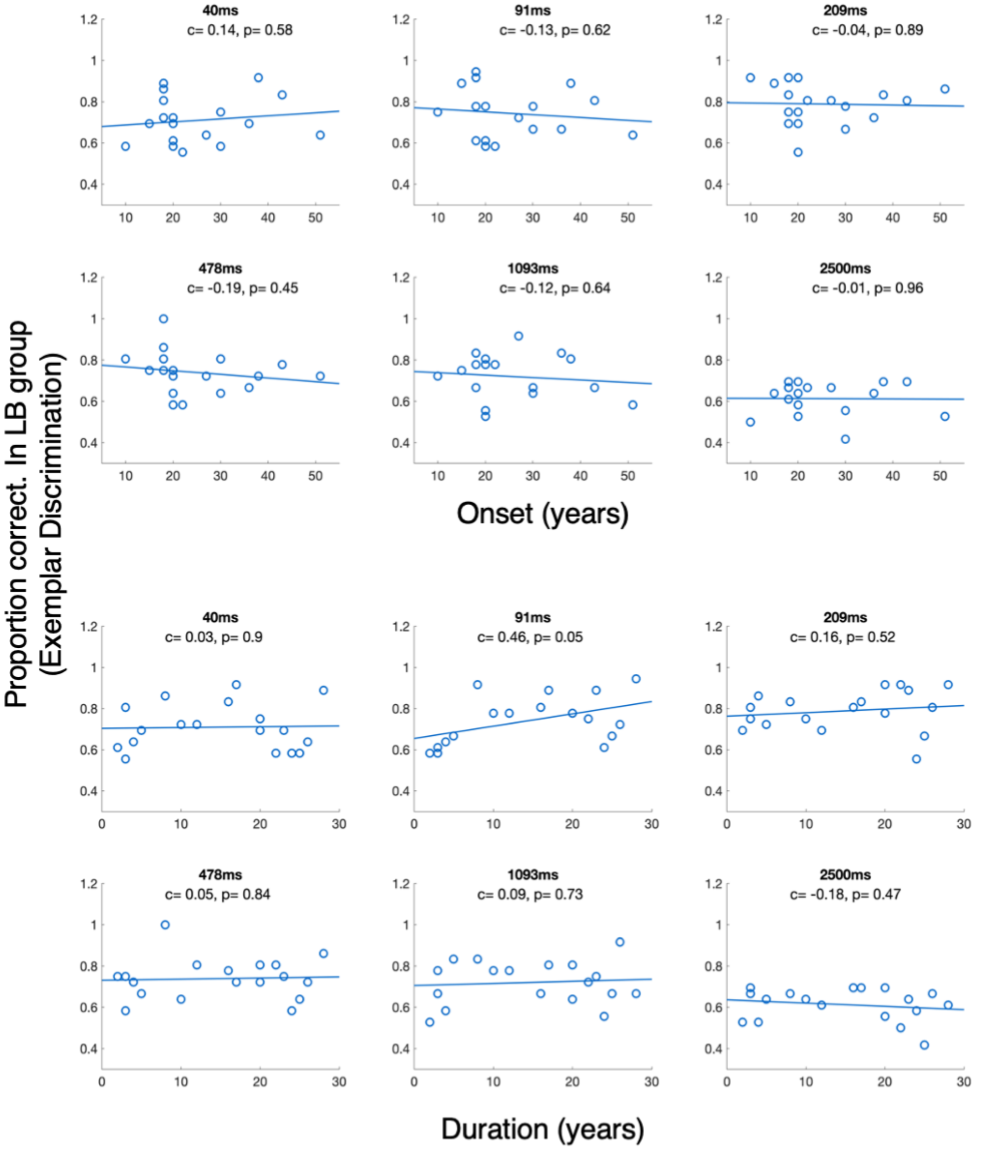

**D**

Correlation Performance Texture with Onset and Duration of Blindness

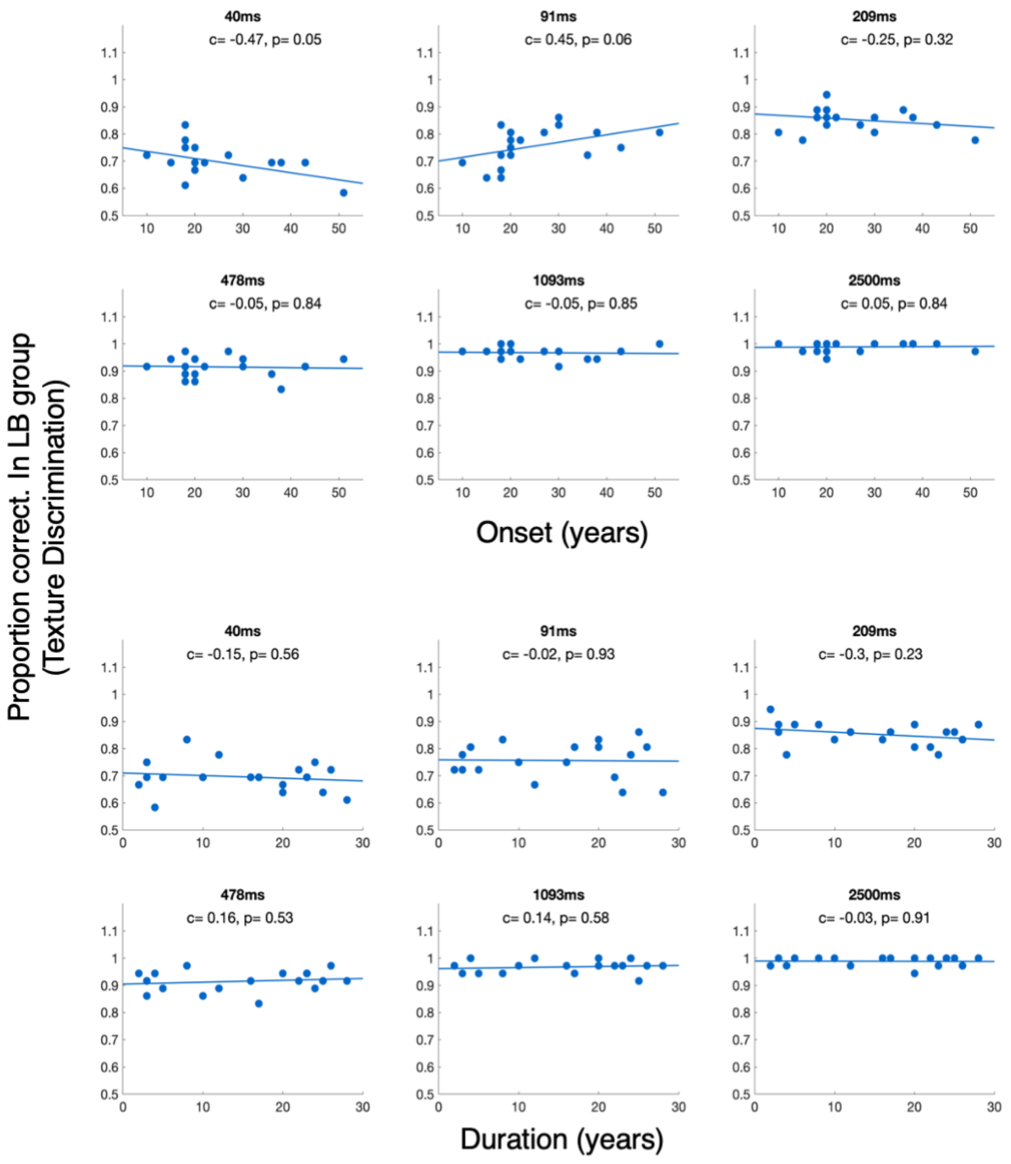


**Figure S3. Correlation between LB group's performance and several variables of interest**
(A, B) Correlation between overall performance of LB in both experiments and participants' onset and duration of blindness (in years).
(C, D) Correlation between performance for LB calculated separately for each duration in Exemplar Discrimination and Texture Discrimination with onset and duration of blindness.
Performance is assessed as proportion of correct answers at Group level, for each experimental condition.

| **Textures 1: used in both experiments** | **Textures 2: only in Texture Discrimination** |
|---|---|
| Applause large crowd | Applause_big_room |
| Applause2 | Applause1 |
| Bathroom sink | Bath being drawn |

| | |
|---|---|
| Bulldozer | Waterfall |
| Castanets1 | Castanets2 |
| Electric adding machine | Teletype city room |
| Fast running river | River running over shallows |
| Fire burning room2 | Fire3 |
| Fire forest inferno | Fire3 |
| Frogs4 | Frogs3 |
| Frying bacon | Crunching cellophane |
| Heavy rain falling and dripping | Heavy rain on hard surface |
| Heavy rain on hard surface | Rain in woods2 |
| Horse trotting on cobblestones | Horse and buggy |
| Industrial machinery | Construction site ambience |
| Jungle rain | Rain in woods1 |
| Linotype | Teletype city room |
| Motorcycle idling | Idling boat |
| Pneumatic drills | Construction site ambience |
| Printing press | Construction site ambience |
| Radiostatic2 | Radio static1 |
| Rain in woods1 | Frogs1 |
| Rain in woods2 | Jungle rain |
| Rain | Rain in woods1 |
| Rhythmic applause | Applause big room |
| River running over shallows | Applause big room |
| Shaking coins | Pouring coins1 |
| Ship anchor being raised | Pneumatic drills |
| Sparrows large excited group | Birds in tropical forest |
| Stream near small waterfall | River running over shallows |
| Teletype city room | Teletype |
| Typewriter IBM electric | Typewriter manual |
| Water running into sink | Bathroom sink |
| Waterfall | Air conditioner |
| Frogs3 | Frogs1 |
| Metal lathe | Blender |
| Applause large crowd | Applause big room |

**Table S1. List of the textures employed in the experiment**
In Texture discrimination, for each texture pair, two excerpts came from exemplars of Textures 1 while the other one came from one excerpt of Textures 2. In Exemplar discrimination only exemplars from Textures 1 were presented (Adapted from McDermott, Schemitsch, and Simoncelli 2013).

1. Late onset blinds group (N=18)

| Participant | Age | Sex | Handiness | Residual Vision | Onset | Etiology | Education | Musical Experience |
|---|---|---|---|---|---|---|---|---|
| ***LB01*** | 21 | M | R | None | 18yo | Congenital Glaucoma | Middle School | No |
| ***LB02*** | 26 | M | R | LP | 18yo | Retinitis pigmentosa | University | No |
| ***LB03*** | 44 | F | R | LP | 20yo | Retinitis pigmentosa | High School (2y) | No |
| ***LB04*** | 25 | F | R | None | 22yo LE14yo RE22yo | Stevens-Johnson syndrome | University | No |
| ***LB05*** | 41 | M | R | None | 36yo | Familial exudative vitreoretinopathy | High School | No |

| | | | | | | | | |
|---|---|---|---|---|---|---|---|---|
| ***LB06*** | 30 | F | R | None | 20yo | Eye tumor | High School | No |
| ***LB07*** | 38 | F | R | None | 15yo | Glaucoma | University | No |
| ***LB08*** | 59 | M | R | None | 43yo | Optic Nerve Sheath Meningioma | Middle School | No |
| ***LB09*** | 55 | F | R | LP | 30yo | Retinitis pigmentosa | University | No |
| ***LB10*** | 52 | F | R | None | 27yo | Retinitis pigmentosa | High School | No |
| ***LB11*** | 55 | F | R | LP, SP, MP | 51yo | Bietti's Crystalline Dystrophy | University | No |
| ***LB12*** | 46 | M | R | LP | 18yo | 6mo: Removed congenital cataract; then Glaucoma | Middle School | No |
| ***LB13*** | 50 | F | R | LP | 11yo | Retinitis pigmentosa | High School | No |
| ***LB14*** | 55 | M | R | LP | 38yo | Retinitis pigmentosa | Middle School | No |
| ***LB15*** | 40 | F | R | None | 20yo | glaucoma | High School | No |
| ***LB16*** | 32 | M | R | LP | 10yo | Optic nerve compression Astrocytoma | University | No |
| ***LB17*** | 22 | M | R | None | 20yo | Glaucoma | High School | No |
| ***LB18*** | 30 | M | R | LP (only RE) | 18yo | Glaucoma (RE) and Retinal detachment (LE) | High School | No |

2. Congenitally blind group (N=18)

| | | | | | | | | |
|---|---|---|---|---|---|---|---|---|
| ***CB01*** | **32** | **M** | **R** | **LP** | **0** | **Congenital Glaucoma** | **University** | **No** |
| ***CB02*** | 36 | M | R | None | 0 | Retinitis pigmentosa | High School | No |
| ***CB03*** | 44 | M | R | None | 0 | Congenital Glaucoma | High School | No |
| ***CB04*** | 60 | F | R | LP | 0 | Retinopathy of prematurity | High School | Yes |
| ***CB05*** | 45 | M | R | LP | 0 | Congenital Glaucoma | University | No |
| ***CB06*** | 43 | F | R | None | 0 | Retinopathy of prematurity | High School | No |
| ***CB07*** | 28 | F | R | LP | 0 | Microphthalmia | University | No |
| ***CB08*** | 29 | F | L | None | 0 | Retinopathy of prematurity | | No |
| ***CB09*** | 27 | M | R | LP | 0 | Retinopathy of prematurity | High School | Yes |
| ***CB10*** | 41 | M | R | LP | 0 | Retinitis pigmentosa | University | No |
| ***CB11*** | 59 | M | R | None | 0 | Glaucoma | High School | Yes |
| ***CB12*** | 37 | F | R/L | LP | 0 | Congenital cataract | High School | No |
| ***CB13*** | 20 | F | R | None | 0 | Microphthalmia and Aniridia | University | No |
| ***CB14*** | 34 | F | R | LP | 0 | Optic nerve hypoplasia | University | No |
| ***CB15*** | 32 | M | R | LP | 0 | Retinopathy of prematurity | University | Yes |
| ***CB16*** | 30 | M | R | LP | 0 | Leber's congenital amaurosis | University | Yes |
| ***CB17*** | 44 | F | R | None | 0 | Virus during pregnacy | High School | No |

| | | | | | | | | |
|---|---|---|---|---|---|---|---|---|
| ***CB18*** | 28 | F | R | None | 0 | Retinopathy of prematurity | University | No |

**Table S2. Characteristics of blind participants**
LP = light perception; SP= Silhouette perception; MP = motion perception; mo = months old; yo = years old; M = male; F = female; LE = left eye; RE= right eye; Musical experience: Yes = professional musical training, studied music for at least 10 years
